# Origin, divergence and migration routes of psyllids of the *Cacopsylla pruni* complex (Hemiptera: Psylloidea) inferred by Approximate Bayesian Computation methods

**DOI:** 10.1101/2024.07.08.602570

**Authors:** Margaux Darnis, Virginie Ravigné, Nicolas Sauvion

**Affiliations:** PHIM, Univ. Montpellier, INRAE, CIRAD, Institut Agro, IRD, Montpellier, France; INRAE, Pathologie Végétale, F-84140, Montfavet, France; CIRAD, UMR PHIM, F-34398 Montpellier, France

**Keywords:** jumping plantlice, population genetics, population divergence, vector-born disease, phylogeography, ABC

## Abstract

Population genetics is essential to decipher the evolutionary history of pests and insect vectors from both a theoretical point of view and to predict and mitigate the future of epidemics. We attempt to shed light on the evolutionary history and phylogeography of two cryptic psyllid species (namely A and B) of the *Cacopsylla pruni* complex, vectors of ‘C*andidatus* phytoplasma prunorum’. It is known to cause the European stone fruit yellows (ESFY), a disease affecting *Prunus* trees and causing significant crop losses each year. Analyses were conducted to decipher the origin, order and time of divergence, as well as the migration routes of the species complex in the Western Palearctic. We used mitochondrial and nuclear gene data to infer the population genetic diversity and structure of the complex, and to reconstruct a dated phylogenetic tree of the Psyllinae family in order to subsequently perform Approximate Bayesian Computation analyses. Both species have diverged in what is now France from a common ancestor, around 20.19 Mya, before expanding into Spain around 6.61 Mya for species A and Eastern Europe around 6.36 Mya for species B. Then species B seem to have passed into Corsica during the Messinian salinity crisis (5.96-5.33 Mya) from French or Italian B populations. No apparent admixture was found between both species after their divergence, which would indicate an absence of gene flow between them at the point when they recolonised common ecological niches. This strong genetic differentiation confirms previous work on reproductive barriers between the two species.

## 1 - Introduction

The great diversity of contemporary insects is the result of a long evolutionary history, strongly influenced by the progressive diversification of terrestrial plants, particularly in phytophagous species (Labandeira 2006; Schatz et al. 2017). The geographical distribution of insects and their host plants, as well as their variations, are a legacy of a distant past marked by major geological and climatic events that have shaped the earth’s surface (Kergoat et al. 2017). Over the past 20 years, biogeography, which aims at understanding the role of geological history, evolution and ecology in the geography and diversification of living organisms, has benefited from considerable methodological and conceptual advances (e.g., Morlon 2014; Maliet et al. 2019; Louca & Pennell 2020; Morlon et al. 2022; Horn et al. 2022; Kawahara et al. 2023). Thanks to molecular phylogeny, for example, it is now possible to infer the respective roles of vicariance (i.e., allopatric speciation resulting from a new geological barrier from within the species range) and dispersal (i.e., the spread of organisms to a new area) in explaining the disjointed distribution of two closely related taxa (e.g., Bossert et al. 2021), and even to propose complex biogeographical scenarios based on alternating cycles of these two processes (Ronquist & Sanmartin 2011).

One of the great difficulties of biogeography is to untangle the web of spatial, temporal and environmental heterogeneity scales on which processes have operated. On the spatial scale of a continent such as the Western Palearctic, the establishment of contemporary flora and fauna has been shaped since the Late Tertiary (i.e., the Neogene: 23.030-2.588 Mya) by climatic upheavals (i.e., warming followed by cooling and drying), and influenced by tectonic phenomena (i.e., northward movement of the African plate leading to the uplift of the young mountains of southern Europe such as the Alps (Mosbrugger et al. 2005; Ezquerro et al. 2022; van Hinsbergen et al. 2020)). These phenomena may have caused major geographic shifts in species distribution and rearrangements in the assemblages of contemporary species or ancestors of present-day species. These phenomena reached their peak during the Pleistocene (2.588-0.0117 Mya) as a result of strong and repeated climatic oscillations between glacial and inter-glacial periods (Hughes & Gibbard, 2018). On the European continent, numerous phylogeography studies have shown the importance of the Iberian Peninsula, Italy and the Balkans as refuge areas for various species of animals and plants during the Ice Age, with expansion towards more northerly regions during periods of warming, with mountains such as the Alps and the Pyrenees playing a major barrier role (Taberlet et al. 1998; Avise 2009; Wallis et al. 2016). While these scenarios of postglacial south-north recolonisation are commonly accepted to explain allopatric speciation or sub-speciation at the European scale, so-called “extra-Mediterranean” refuges (i.e., refuges in Central Europe) have also been put forward to explain the observed genetic structure of present-day populations in Europe (Stewart & Lister, 2001; Schmitt & Varga, 2012; Nokkala et al. 2022). In addition, in several cases, lineages that have been geographically isolated over geological time have met again as they expanded, forming hybrid zones where introgression has left complex genomic signatures (Taberlet et al. 1998; Hewitt 2001; Taylor et al. 2015). These different patterns of colonisation, by vicariance and/or dispersion, at various spatial and temporal scales, and the possibility of encounters between isolated lineages show the extent to which the evolutionary history of even closely related species can be the result of a complex natural scenario. In the specific case of insect pests or vectors, another complexity factor can be added: human activity, particularly agriculture. Human-induced movements can influence the distribution and demography of populations and greatly accelerate their expansion (Liebhold & Tobin 2008). These factors can change population genetic structure and add new genetic signatures on already existing natural biogeographical patterns (Stone et al. 2007).

To decipher scenarios with complex biogeographic patterns, the Approximate Bayesian Computation (ABC) method is an increasingly used method in population genetics for statistical inference (Beaumont 2002; Csilléry et al. 2010; Sunnåker et al. 2013). Among other things, it has been used to reconstruct species divergence and migration scenarios (Fresia, Azeredo-Espin, & Lyra 2013; Sajidet al. 2014), invasion scenarios (Zepeda-Paulo et al. 2010; Ascunce et al. 2011; Barrès et al. 2012; Mallez et al. 2021), scenarios of glacial refugia and postglacial recolonisation (Bidegaray-Batista et al. 2016; Sim et al. 2016; Song et al. 2018), as well as habitat colonisation scenarios (Estoup et al. 2001; 2004; Thornton & Andolfatto 2006; Pascual et al. 2007). In ABC, the limiting step of very complex likelihood computations is replaced by an approximation of the likelihood, which is obtained by simulating datasets under considered scenarios and selecting the simulated datasets that are closest to the observed data using a regressive approach on summary statistics (Beaumont 2002).

The aim of this study was to decipher the evolutionary history of the psyllid complex *Cacopsylla (Thamnopsylla) pruni* (Scopoli 1763) [Hemiptera: Psylloidea] in Europe. These insects are known to be the vectors of the bacterium ‘*Candidatus* phytoplasma prunorum’, a phytopathogen responsible for the European stone fruit yellows (ESFY) a disease affecting fruit trees of the *Prunus* genus (Lee et al. 1998; Carraro et al. 1998; Danet et al. 2011; Marie-Jeanne et al. 2020). *Cacopsylla pruni*, first thought to be a single species (Carraro et al. 1998), was recently proven to be a complex of two cryptic species (Peccoud et. 2013; Peccoud et. 2018; Sauvion et al. 2024), hereafter referred to as A and B. The two species are nearly impossible to distinguish on the basis of morphological criteria alone, but their level of genetic differentiation was shown to be very strong in relation to an extreme reproductive isolation: *in natura*, inter-species mating has been observed but hybrids are extremely rare (< 1/1000) (Peccoud et al. 2018). The two species are present in sympatry or strict allopatry, depending on the geographic area, and they are endemic to the wild habitats of the Western Palearctic (Sauvion et al. 2021). Ecologically, both species share the same biological cycle and both can spread the pathogen (Marie-Jeanne et al. 2020). They both have an annual life cycle. They breed in spring on wild *Prunus* trees (e.g., blackthorn, *Prunus spinosa* L.) and take refuge on conifers (i.e., shelter plants) for the rest of the year, probably to escape the heat of summer (Thébaud et al. 2009). These species are therefore capable of dispersing over great distances (at least 50 km) and adapting to different ecological niches (mid-altitude forests vs. lowland shrubs) (Marie-Jeanne et al. 2020).

The sympatric occurrence of the two cryptic species, in addition to their unique distribution pattern [e.g., only species B is found in Corsica, an island off of the coast of southern France, raises the question of their evolutionary history, given that the French coast facing the island shows sympatry between both species, seemingly indicating that psyllids are not capable of crossing the sea (Sauvion et al. 2021)]. In the present study, we focussed on the time and mode of both the divergence and expansion of the two species of the *C. pruni* complex in the Western Palearctic. We used mitochondrial and nuclear DNA sequences to generate a phylogenetic tree of a part of the Psyllinae family comprising the *Cacopsylla* genus, and to describe the genetic diversity and structure of the *C. pruni* complex in Europe in order to reconstruct hypotheses of its evolutionary history. We then used an ABC approach to compare competing evolutionary scenarios and to attempt to identify recent and old evolutionary events that shaped the current population genetic structure of these insect vectors.

## 2 - Materials and Methods

### 2.1 Samples and genetic markers

For the study, 97 sample sites (i.e., geolocated sampling points named after the nearest locality), representing a total of 1245 individuals from all across Europe, were analysed: 71 sites from France, six sites from Spain, and 20 sites from Central and Eastern Europe. The samples are listed and described in Table S1 and their geographical locations are shown in Fig. S1. Sampling was performed between 2005 and 2018. For all the following analyses, we considered three geographical areas: species A allopatry area (Spain = S), species B allopatry area (Central and Eastern Europe = CEE), and species A and B sympatry area (France = F). For the ABC analyses, we also considered Corsica, a B allopatry area.

Two genetic markers were used: 621 bp of the mitochondrial DNA (mtDNA) region Cytochrome c oxidase subunit I -t-tRNA-COII, referred to as COI-t in the rest of the study, obtained with the UEA9 and C2-N-3389 primers, as described in Percy (2003) (213 sequences of species A, and 331 sequences of species B were used for ‘Descriptive genetic analysis’ (see below)), and 753 bp of the nuclear (ribosomic) DNA region Internal transcribed Spacer 2 (ITS2), obtained with the CAS5p8sFcm and CAS28sB1d primers designed by Peccoud et al. (2013) (340 species A and 443 species B sequences). The choice of this mitochondrial region made it possible to use the sequences referred to in the article by Percy (2003) that were available in GenBank. Sanger genotyping was carried out by the French National Sequencing Centre, Genoscope. The sequences were analysed with Geneious Pro 10.2.6 according to the following steps: for each individual, the forward and reverse sequences were aligned, compared and assembled to obtain a consensus sequence. Particular attention was paid to the mitochondrial sequences because it is known that nuclear copies of mitochondrial genes can generate ambiguous results (Hurst & Jiggins 2005; Song et al. 2008). Every double pic was visualised to ensure that the haplotypes were functional coding genes (no indels or stop codons). Finally, each individual was assigned to the A or B species by following the diagnostic PCR protocol described by Peccoud et al. (2013).

For the phylogenetic analysis, we analysed 368 sequences of the COI-t-tRNA-COII region of the mitochondrial genome, including 323 sequences (92 species) of Psyllinae, 217 (37 species) which were of the *Cacopsylla* genus. A total of 232 sequences (103 species) were extracted from Genbank, including 66 sequences (34 species) available from the articles published by Percy (2003a, 2003b) and Percy et al. (2018). The other 136 sequences (21 species) were obtained from our own samples. A preliminary analysis allowed us to identify the sequences correctly aligning with ours and the mislabelled sequences (not shown). After this step, 94 sequences were kept for our study and the others were discarded. The table of the sequences we used, their origins and GenBank numbers are available in Table S2.

### 2.2 Phylogenetic analyses and molecular dating

The Psylloidea classification published by Percy et al. (2018) based on the almost complete mitochondrial genome of 359 species of Psylloidea was used as a reference for our tree. This classification was reassuringly congruent with earlier classifications based on morphological markers and also with later classifications based on molecular markers published by Cho et al. (2019) and Burckhardt et al. (2021) for the taxa of interest (*Cacopsylla* genus, Psyllinae family) (Percy 2003a; Percy et al. 2018; Cho et al. 2019; Burckhardt et al. 2021). These phylogenies constitute the current consensual hypothesis on the phylogeny of Psylloidea.

Nucleotide sequence alignment was performed with the MAFFT program (multiple sequence alignment) and the sequences were cleaned with the BMGE (Block Mapping and Gathering with Entropy) software (Criscuolo & Gribaldo 2010; Katoh & Standley 2013). The phylogenetic tree of the part of the Psyllinae family comprising the *Cacopsylla* genus was reconstructed using Maximum likelihood (ML) with the PhyML method (Guindon & Gascuel 2003; Guindon et al. 2010). The resulting tree was visualised with a Newick display (Junier & Zdobnov 2010). All these steps were conducted with NGPhylogeny, an online platform that embeds a large range of tools and methods for phylogenetic analysis (Lemoine et al. 2019). Approximate Bayes (aBayes) branch support and bootstrap values were also calculated with NGPhylogeny as it was shown that aBayes, a parametric branch support method included in PhyML, offers the highest statistical power, whereas non-parametric tests like classic bootstrap were shown to be much less powerful but more (excessively) conservative (Maria et al. 2011). Finally, the phylogenetic tree was formatted with the Interaction Tree Of Life (iTOL) v5 program (Letunic et Bork 2021) and it was verified that the tree obtained with this method was congruent with the one obtained by Percy et al. (2018). Tree inferences were also carried out with the FastME (distance-based) method implemented in NGphylogeny and with a ML method implemented in MEGA11 v11.0.13 (Hall 2013; Mello 2018) for comparison purposes. To infer the date of divergence between the two species of the *C. pruni* complex, a molecular dating analysis was performed using MEGA11 v11.0.13 (Mello 2018). To compute the timetree in MEGA11, the RelTime-ML method was chosen as it is the original method proposed by Tamura et al. (2012) and is suitable for most data. Within this option, time estimates are computed based on branch lengths and optimised by maximum likelihood (ML), which is a robust statistical method (Tamura et al. 2012). The BMGE aligned and cleaned sequences and phylogenetic tree inferred with PhyML in NGphylogeny were uploaded to MEGA11. The outgroup for the phylogenetic analysis was chosen to be the *Cyamophila* clade [*Cyamophila floribundae* Cho & Burckhardt, 2017, *Cyamophila prohaskai* (Priesner, 1927) and *Cyamophila willieti* (Wu, 1932)], a sister taxon to the *Cacopsylla* clade, the clade of interest, according to the classification made by Percy (2003a) and Percy et al. (2018), and by others since then (Cho et al. 2019; Burckhardt et al. 2021). Analysis settings were established as follows: the GTR (General Time Reversible model) + G + I (gamma distributed with invariant site rates among sites) was chosen as the substitution model that was used to estimate branch lengths. To make this choice, we used the feature proposed by MEGA11 that chooses the best model based on a comparison of all the substitution models available and the BIC, AICc and lnL criteria (Table S3: comparison between substitution models) (Hall 2013). The *Arytinnis* MRCA was calibrated using the age of the Canary Island formation (16 Mya) as an upper bound as it was shown that the *Arytinnis* clade is almost exclusively composed of endemic Canary Island species and suggested that the basal node of this clade was dated to after the formation of the islands (Percy 2003a). The *Cacopsylla* MRCA was calibrated using (i) fossil data (Becker 1864; Klimaszewski 1993) for its lower bound based on the age of the oldest *Cacopsylla* fossil we could identify from the literature, and (ii) the date of divergence between psyllids and aphids (Johnson et al. 2018) as an extreme upper bound. The five other nodes were calibrated using dated host phylogenies. Multiple studies have shown that Psylloidea have a very narrow host range corresponding to low-level plant taxa, either a specific plant family or genus, that they often have a one-to-one relationship with a host plant at the species level, and that they are usually monophagous (Hodkinson 1974; 1980, Burckhardt & Lauterer 1989; Percy 2003a; Hodkinson 2009; Burckhardt et al. 2014; Ouvrard et al. 2015). Congruently, when analysing the Psyllinae phylogenetic tree obtained in the present study, we observed considerable congruence between the clustering of the psyllid species with their Rosaceae host plant phylogeny (Fig. S2). These non-random associations have already been reported in other studies (Percy et al. 2004; Percy et al. 2018; Burckhardt et al. 2021). These observations and the prevalence of highly specific host-plant associations in psyllids have caused a number of authors to consider a possible co-evolution or co-speciation between psyllids and their hosts (Hodkinson 1984; White & Hodkinson 1985; Brown & Hodkinson 1988; Van Klinken 2000; Percy et al. 2004). We have retained this hypothesis as the base assumption to calibrate some nodes, even if we cannot exclude the hypothesis of a host switch that would not have been contemporaneous to host divergence (Percy et al. 2004; Ouvrard et al. 2015). The nodes were thus calibrated with the lower minimum and upper maximum values of the confidence intervals estimated for their equivalents in the Rosaceae dated phylogenetic trees (Chin et al. 2014; Zhang et al. 2017; Korotkova et al. 2018; Du et al. 2021). All the calibration points, their prior distribution and the corresponding references are shown in Fig. S2.

### 2.3 Descriptive genetic analyses

For the mitochondrial (COI-t) and nuclear (ITS2) genes, the length of the sequences, the number of sequences, sites, identical sites, polymorphic sites (S), the total number of mutations, number of haplotypes (H), haplotype diversity (Hd), nucleotide diversity (Pi), Tajima’s D and its P-value were calculated using DnaSP 6.12.3 (Rozas et al. 2017). For the COI-t gene, the same indices were computed for each geographical area studied, and an analysis of the divergence between geographical areas was conducted for both species. Those supplementary analyses were not conducted on the ITS2 gene since we mainly had only French samples for this gene. Haplotype networks of both genes were generated by the Median Joining Network method of PopART 1.7 software (Leigh & Bryant 2015), an appropriate method for network reconstruction of intraspecific data (Bandelt et al. 1999; Woolley et al. 2008). PopART was also used to visualise the proportion of each haplotype in each sample site for both genes and species with pie charts. All these analyses were performed independently on each species. The number of private haplotypes (i.e., haplotypes found in only a given geographic area) of the A and B species was computed for the COI-t gene by a rarefaction approach using HP-rare software (Kalinowski 2005). Rarefaction is a method initially used to compute allelic richness for samples of different sizes, i.e., accounting for the probability of sampling a higher number of alleles in larger samples, and ultimately allows us to obtain comparable statistics unbiased by sample size (Kalinowski 2004; Sjöstrand et al. 2014).

### 2.4 ABC analyses

To explore the possible evolutionary scenarios of the *C. pruni* complex, an ABC (Approximate Bayesian Computation) analysis was performed on the data of the COI-t gene using DIYABC 2.1 software (Cornuet et al. 2008; 2014). It was used to compare several competing hypotheses on demographic history, population divergence and expansion of *C. pruni* populations at the Western Palearctic scale. To do so, the software simulates a huge number of datasets by picking different parameter values from the prior distributions chosen by the user under several scenarios also given by the user. The program then estimates the posterior probability of each competing scenario and infers the a posteriori demographic parameter values and distributions under a given scenario by comparing the simulated and observed datasets on chosen summary statistics (Cornuet et al. 2008).

In the present analyses, we were interested in the evolutionary history of the two species A and B and their distribution along the three geographical areas (S, CEE and F), leading us to consider four “populations” to simulate in ABC analyses: A-S, A-F, B-F and B-CEE. However, following the methodological and conceptual guidelines recommended by Lombaert et al. (2014), we were careful not to pool samples from different sites by geographical area, but instead used them as replicates of the same evolutionary history in the ABC analyses. Indeed, even if pooling is common practice, it can give misleading results and lead to misinterpretation of the ABC analyses (Lombaert et al. 2014). Thus, several sample sites were chosen for each geographical area to be used as replicates for ABC analyses. They were chosen on the basis of three criteria: (i) geographical distribution: we chose sites not too close to each other to best represent the true distribution of the species in the area; (ii) number of sequences: sites with the largest number of sequenced individuals were privileged; (iii) availability of data: while ten sites were retained in CEE and F, only six sites were available for Spain, two of which were very close to each other, so only five sites were retained for this area.

The hypotheses regarding population divergence, expansion and demographic history were constructed primarily to test the order and time of divergence of population groups, the geographical area of the origin population, and the possibility of admixture between these population groups after divergence. Tests were carried out hierarchically (Fig. 1). First, we tested five competing scenarios of population divergence without admixture, assuming that the date of divergence between species A and B was the one inferred by our phylogenetic and molecular dating analysis (step A). We then tested two competing scenarios of population admixture after divergence between species A and B, including one with no admixture (step B). Using the results of the last step, we tested four competing scenarios of population admixture within each species, once again including one scenario with no admixture (step C). Finally, we tested scenarios for the divergence of the Corsican population. We tested six scenarios for divergence of the Corsican population: before/during separation from the continent (15-28 Mya), during the Messinian salinity crisis (5.33-5.96 Mya), and before/during separation from the continent with admixture during the Messinian salinity crisis. The first three scenarios infer divergence from French populations and the last three infer divergence from the CEE populations, in this case most probably the Italian ones (step D). In each case, we simulated a total (number or scenario) x 10^6^ datasets for each scenario in order to obtain both computationally and statistically robust results, and used the logistic regression approach implemented in DIYABC to estimate the posterior probabilities of each scenario using 1% of the datasets closest to the observed data. In total, 62 summary statistics were used for the analyses. The final prior definitions and distributions are provided in Table 1 and the summary statistics used for the analyses are provided in Table S4.

**Fig. 1:**
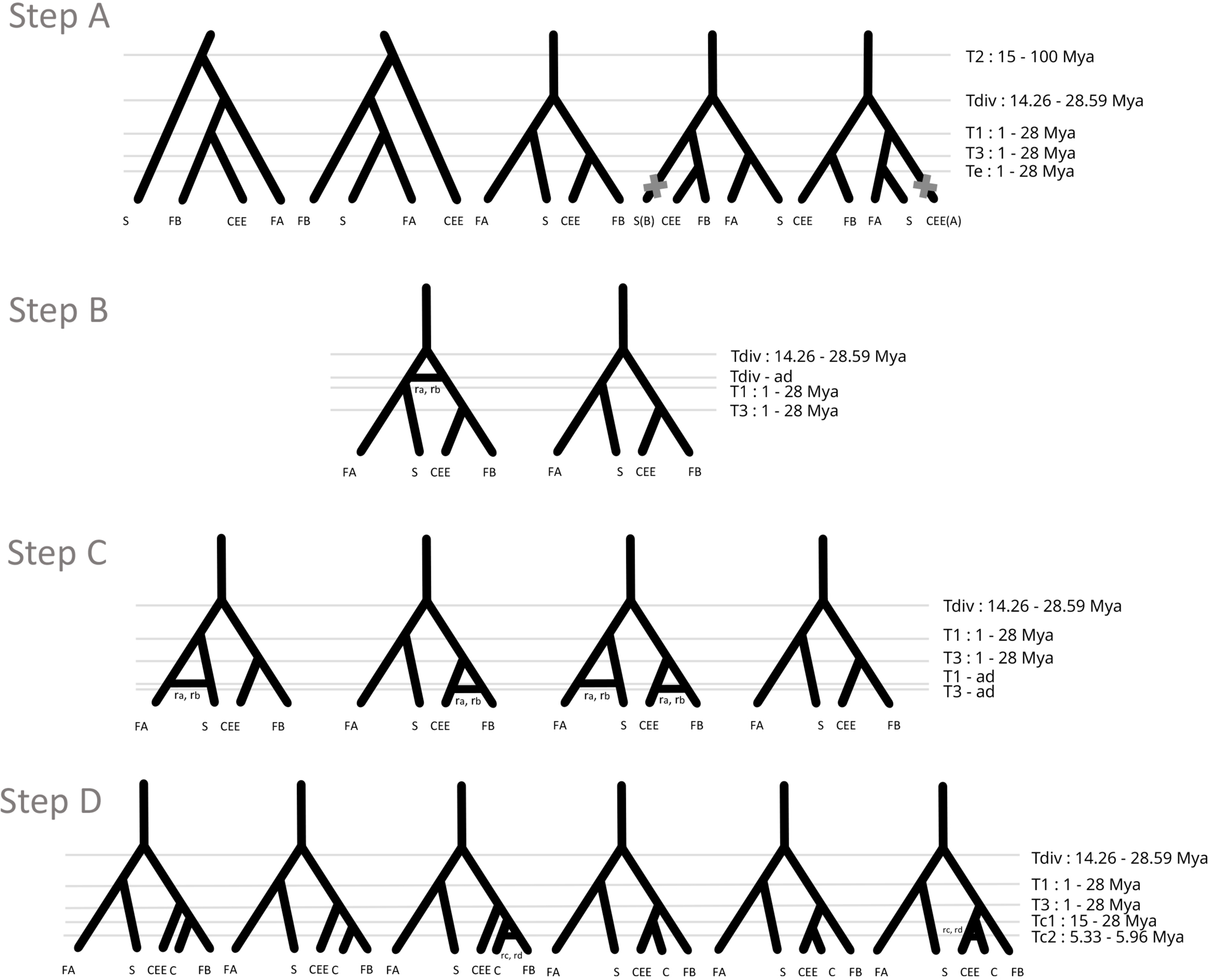
Graphical representation of the scenario tested in each step of the ABC analyses. FA = French populations of species A; FB = French populations of species B; S = Spanish populations; CEE = Central and Eastern European populations; Tdiv = Time of divergence between A and B species; T2 = Time of divergence between origin population and French populations (Sc1 & 2); T1, T3, Te = Time of divergence between populations; Tc1 = Time of divergence of the Corsican population (28-15 Mya); Tc2 = Time of divergence between populations (5.96-5.33 Mya); ad = time of admixture after divergence; ra,rb,rc,rd = admixture rates.

**Table 1:**
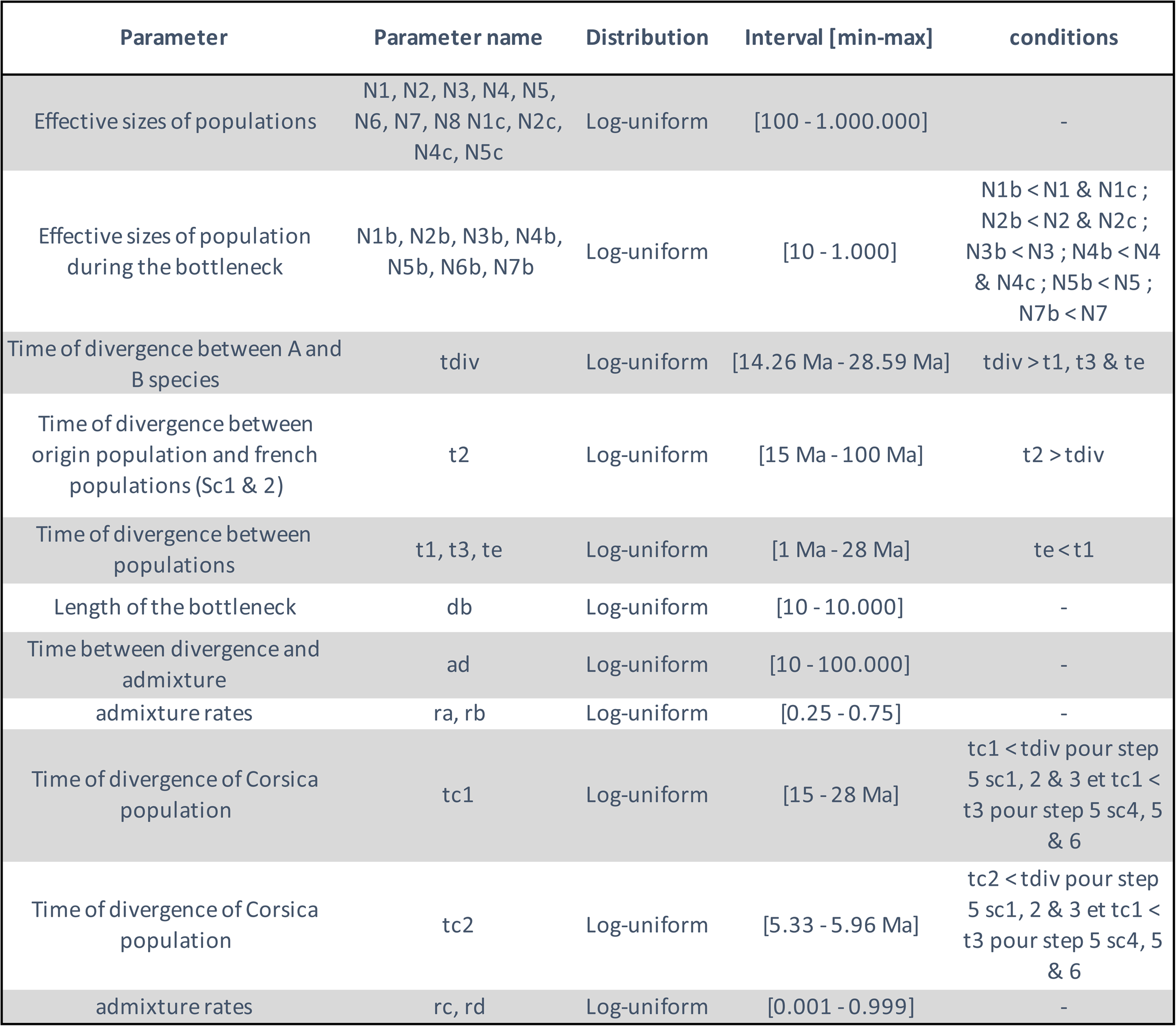
Prior definition and distribution used for the ABC analyses. For each prior used in the ABC analysis, its name, abbreviation, the distribution and minimum/maximum interval of its values and the conditions we set for it are given.

| Parameter | Parameter name | Distribution | Interval [min-max] | conditions |
| --- | --- | --- | --- | --- |
| Effective sizes of populations | N1, N2, N3, N4, N5, N6, N7, N8 N1c, N2c, N4c, N5c | Log-uniform | [100 - 1.000.000] | - |
| Effective sizes of population during the bottleneck | N1b, N2b, N3b, N4b, N5b, N6b, N7b | Log-uniform | [10 - 1.000] | N1b < N1 & N1c ;<br>N2b < N2 & N2c ;<br>N3b < N3 ; N4b < N4 & N4c ; N5b < N5 ;<br>N7b < N7 |
| Time of divergence between A and B species | tdiv | Log-uniform | [14.26 Ma - 28.59 Ma] | tdiv > t1, t3 & te |
| Time of divergence between origin population and french populations (Sc1 & 2) | t2 | Log-uniform | [15 Ma - 100 Ma] | t2 > tdiv |
| Time of divergence between populations | t1, t3, te | Log-uniform | [1 Ma - 28 Ma] | te < t1 |
| Length of the bottleneck | db | Log-uniform | [10 - 10.000] | - |
| Time between divergence and admixture | ad | Log-uniform | [10 - 100.000] | - |
| admixture rates | ra, rb | Log-uniform | [0.25 - 0.75] | - |
| Time of divergence of Corsica population | tc1 | Log-uniform | [15 - 28 Ma] | tc1 < tdiv pour step 5 sc1, 2 & 3 et tc1 < t3 pour step 5 sc4, 5 & 6 |
| Time of divergence of Corsica population | tc2 | Log-uniform | [5.33 - 5.96 Ma] | tc2 < tdiv pour step 5 sc1, 2 & 3 et tc1 < t3 pour step 5 sc4, 5 & 6 |
| admixture rates | rc, rd | Log-uniform | [0.001 - 0.999] | - |

## 3 - Results

### 3.1 Phylogenetic tree of Psyllinae and divergence time estimation of *C. pruni*

Phylogenetic trees were constructed based on the COI-t gene sequences of 94 species. The trees obtained with FastMe (distance based) and ML on MEGA were nearly identical to those of the PhyML analyses made with NGPhylogeny. Thus, only PhyML results are reported here (Figs. 2, S3 and S4). The phylogenetic history for most of the clades was congruent with the consensual hypothesis on the phylogeny of Psylloidea proposed in Percy et al. (2018) and other classifications (Cho et al. 2019; Burckhardt et al. 2021). The *Cacopsylla* clade (in group O of Percy et al. 2018), the largest psyllid genus (∼450 species), was found to be monophyletic with a high support (support value > 0.99), and separated into two clades as in Percy et al. (2018). One clade, essentially psyllids of the *Thamnopsylla* subgenus, was strongly supported (support value > 0.99), and the second, including *Cacopsylla* sensus stricto and *Hepatopsylla*, was moderately supported (support value = 0.81). In addition, *Spanioneura buxi* (Linné) clustered as a sister taxon of *Psylla foersteri* (Percy et al. 2018). Moreover, the present phylogeny comprised a highly-supported (support value > 0.99) clade of taxa comprising the *Arytaina*, *Arytainilla*, *Livilla* and *Arytinnis* clades. The latter three clades clustered together as a sister to the *Arytaina* clade, like in group P of Percy et al. (2018). In addition, the *Arytaina*, *Arytainilla* and *Arytinnis* clades revealed themselves to be monophyletic, whereas *Livilla* turned out to be paraphyletic with respect to *Arytainilla* and *Arytinnis*, as in Burckhardt et al. (2021) and Percy et al. (2003). The *Arytaina* clade was only moderately supported (support value = 0.43), but the *Arytainilla*, the three *Livilla* and the *Arytinnis* clades were all strongly supported (for all, support values = 1, except for the first *Livilla* clade with support value = 0.98). This tree was then used as a backbone for the subsequent molecular dating analysis.

**Fig. 2:**
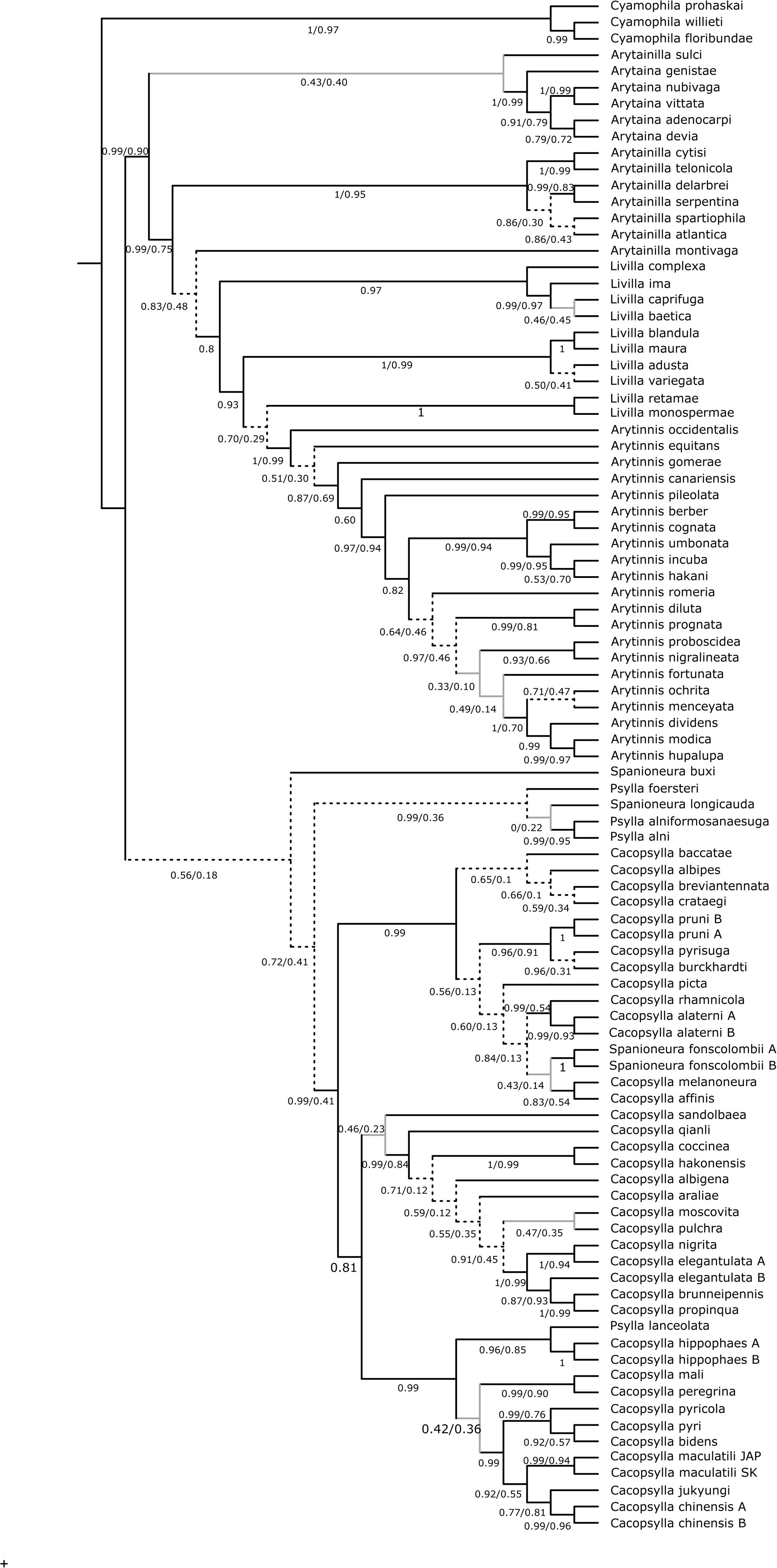
Maximum likelihood phylogenetic tree based on the COI-tRNA-COII mitochondrial gene of 94 Psyllinae species. The number on/under the branches represents the approximate Bayes (aBayes) branch support and bootstrap values of the branches in percentage. Grey branches are branches for which both aBayes and bootstrap are < 50%. Dotted branches are branches for which aBayes is > 50%, but bootstrap is < 50%.

The molecular dating analysis allowed us to estimate the age of the nodes in the phylogenetic tree and to compute a chronogram of the part of the Psyllinae family that contains the *Cacopsylla* genus (Figs. 3 and S5). Divergence time estimates suggested that both *C. pruni* species diverged from a shared common ancestor around 20.19 Mya (95% CI: 14.26-28.59 Mya). This date was hereafter used in the ABC analyses.

**Fig. 3:**
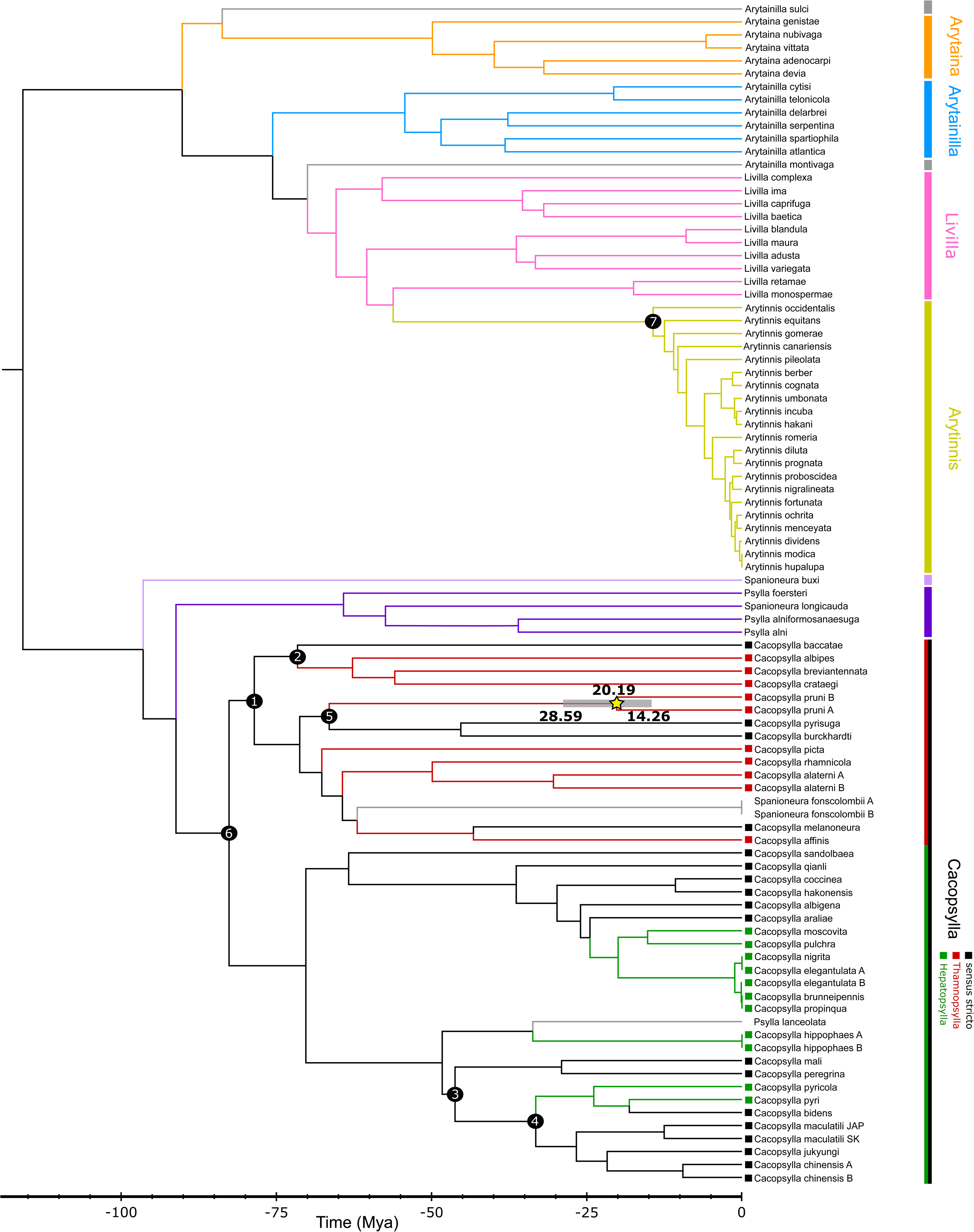
Dated phylogeny (ML) of the Psyllinae based on the COI-tRNA-COII mitochondrial gene of 94 Psyllinae species obtained with MEGA. Black circles represent the seven nodes that were calibrated for the analysis; each colour is associated with a clade. The date represented is the divergence date of both species of the *C. pruni* complex. External group not shown.

### 3.2 Genetic diversity and structure in *C. pruni* regional groups

Table 2 shows the genetic variability of each regional group. For the COI-t gene, the species A populations of France showed higher haplotype and nucleotide diversity than the Spanish ones, and lower haplotype and nucleotide diversity than the Italian and Swiss populations. As for species B, French populations had higher diversity indices than CEE ones.

**Table 2:** Descriptive analysis and statistics of the COI-t and ITS2 genes in each geographical area and total population. Tot = total population; F = France; S = Spain; I+Sw = Italy+Switzerland; CEE = Central and Eastern Europe; bp = base pair, div = diversity

| Group (gene,<br>species &<br>geographical<br>area) | Seq | Length<br>(bp) | Sites | Identical<br>sites | Polymorphic<br>sites (S) | Mutations | Haplotypes<br>(h) | Haplotype<br>div (Hd) | Nucleotide<br>div (Pi) | Parcimony<br>Informative<br>sites | Tajima's<br>D | P-value |
| --- | --- | --- | --- | --- | --- | --- | --- | --- | --- | --- | --- | --- |
| COI-t A tot | 212 | 621 | 581 | 570 | 11 | 11 | 13 | 0.395 | 0.00086 | 4 | -1.72889 | P > 0.05 |
| COI-t A - F | 157 | 621 | 621 | 611 | 10 | 10 | 11 | 0.322 | 0.00064 | 2 | -1.87024 | P < 0.05 |
| COI-t A - S | 39 | 621 | 581 | 577 | 4 | 4 | 5 | 0.242 | 0.00044 | 1 | -1.75782 | 0.1 > P > 0.05 |
| COI-t A - I+Sw | 16 | 621 | 621 | 620 | 1 | 1 | 2 | 0.500 | 0.00081 | 1 | 1.30896 | P > 0.10 |
| COI-t B tot | 332 | 621 | 621 | 590 | 31 | 33 | 34 | 0.347 | 0.00065 | 10 | -2.48958 | P < 0.001 |
| COI-t B - F | 182 | 621 | 621 | 597 | 24 | 24 | 24 | 0.363 | 0.00068 | 3 | -2.49452 | P < 0.001 |
| COI-t B - CEE | 150 | 621 | 621 | 609 | 12 | 13 | 15 | 0.327 | 0.00061 | 7 | -2.14010 | P < 0.05 |
| ITS2 A (F) | 339 | 735 | 735 | 730 | 5 | 5 | 8 | 0.622 | 0.00784 | 4 | 1.31188 | P > 0.10 |
| ITS2 B (F) | 444 | 744 | 744 | 734 | 10 | 11 | 12 | 0.521 | 0.00369 | 10 | 1.4567 | P > 0.10 |

For species A, 13 haplotypes of the COI-t gene were identified among the 212 individual samples, and eight haplotypes of the ITS2 gene for 339 individuals were revealed. In species B, 34 (COI-t, on 332 individuals) and 12 (ITS2, on 444 individuals) haplotypes were revealed. Figures 4 and 5 show haplotype networks and distributions for both genes and both species. Some haplotypes were shared between geographical areas, mainly the most common ones. For each species and each gene, French haplotypes represent the majority of haplotypes identified. This could be due to the high number of French sites sampled. However, after a rarefaction analysis (of the number of private haplotypes per geographical area), accounting for the unequal sample sizes between France and other geographic areas, the observation of a higher haplotype diversity in France is still valid. France has a higher number of private haplotypes than the other geographic areas sampled. With over 100 theoretically sampled individuals per geographical area, species A should show 4.27 private haplotypes in France, 3.47 in Spain and 0.22 in Italy and Switzerland, and species B should show 9.17 private haplotypes in France, 7.98 in Central and Eastern Europe, 2.03 in Corsica and 0 in Spain.

**Fig. 4:**
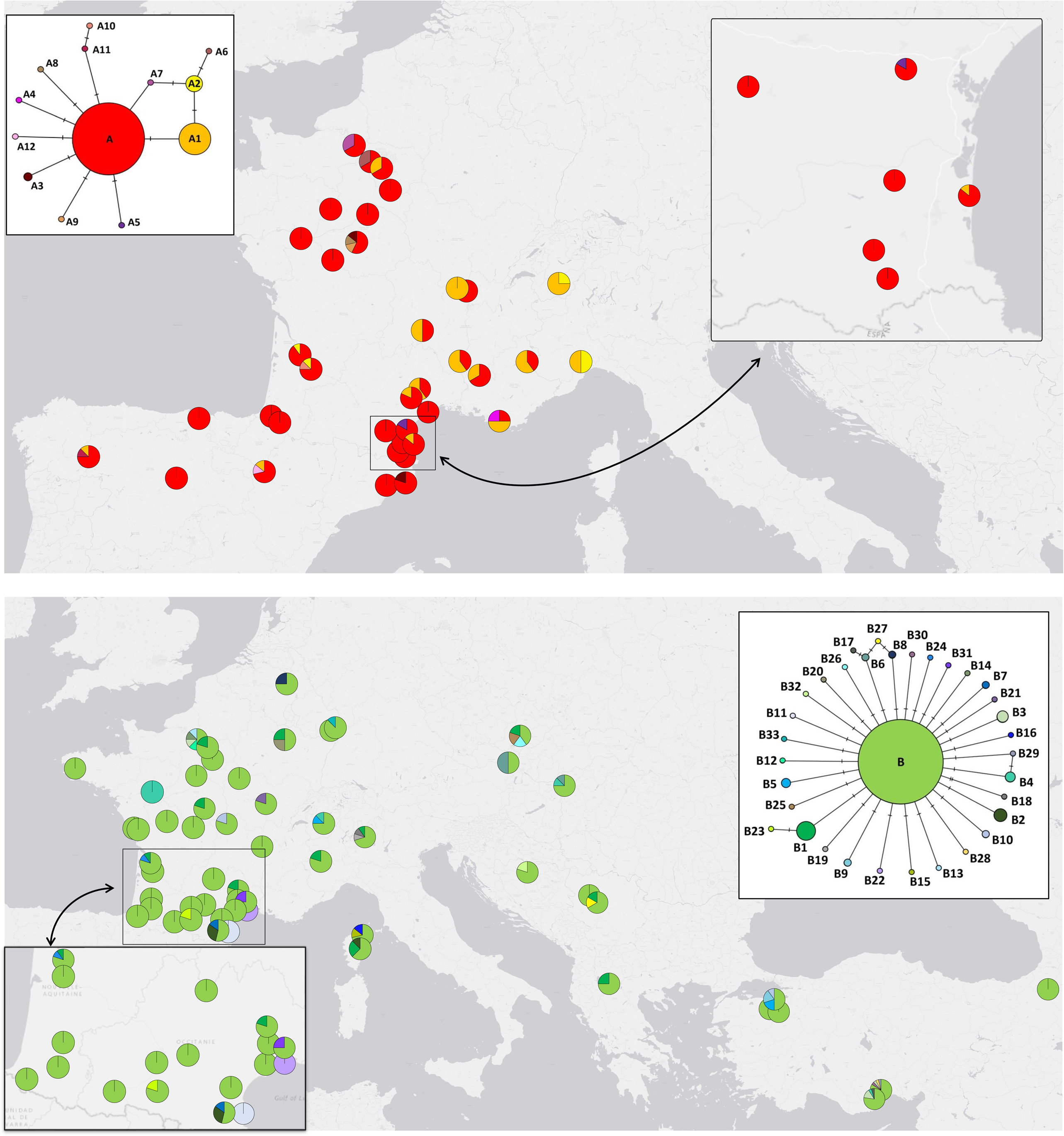
Haplotype network and distribution map for the COI-t gene for both species in the Western Palearctic (species A in red and species B in green) Each colour represents a haplotype. The pie charts represent the proportion of each haplotype in the population.

**Fig. 5:**
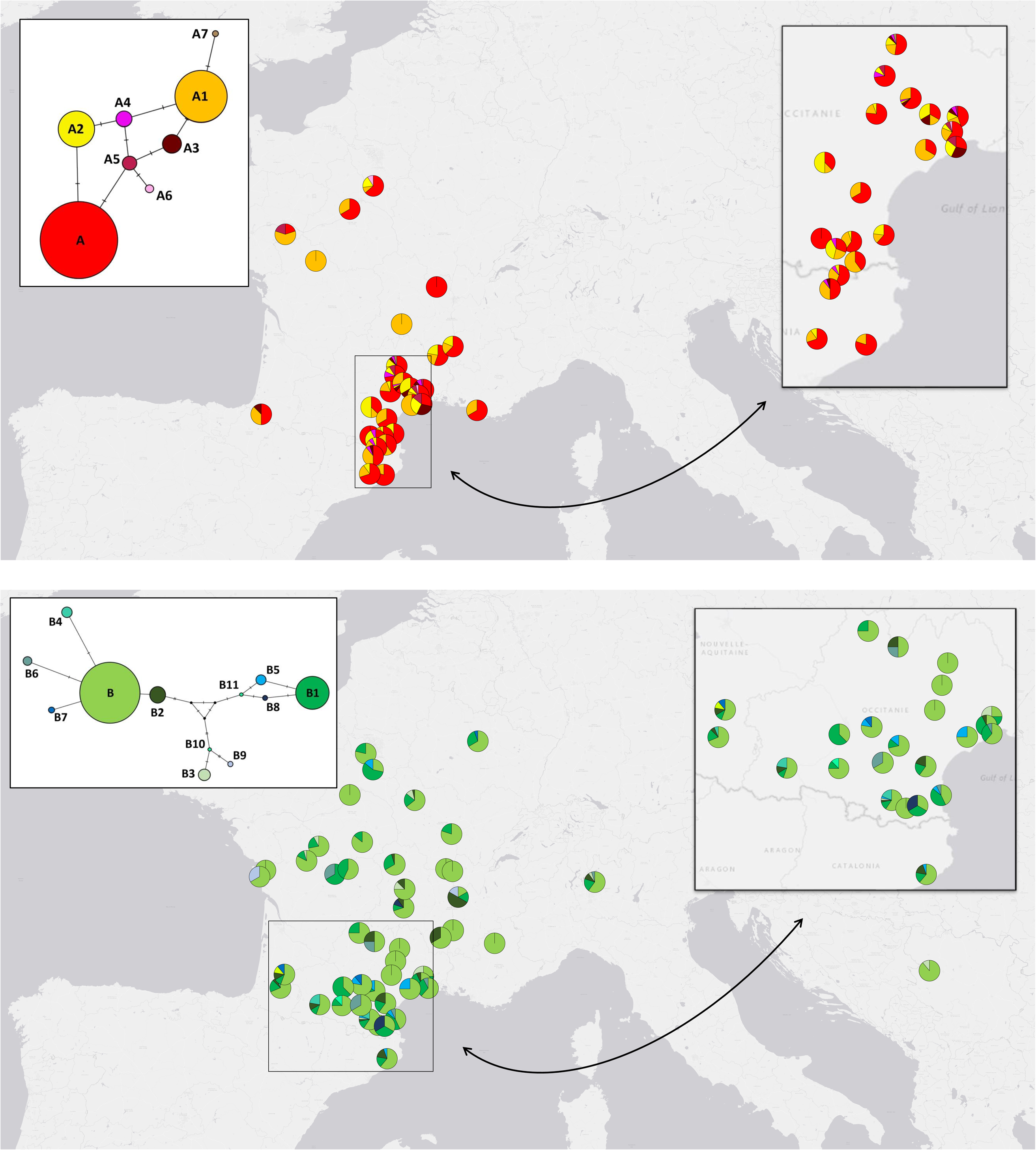
Haplotype network and distribution map for the ITS2 genes for both species in the Western Palearctic (species A in red and species B in green) Each colour represents a haplotype. The pie charts represent the proportion of each haplotype in the population.

In addition, DnaSP made it possible to study the distribution of mutations that were polymorphic in a focal region and monomorphic elsewhere. Such mutations were most frequent in French populations. For species A, seven mutations were polymorphic in France and monomorphic in Spain, whereas only one mutation was polymorphic in Spain and monomorphic in France. Moreover, nine mutations were polymorphic in France and monomorphic in the Italian and Swiss samples, whereas there were no polymorphic mutations in the Italian and Swiss samples and no monomorphic mutations in France. Finally, four mutations were polymorphic in Spain and monomorphic in the Italian and Swiss samples, whereas only one was polymorphic in the Italian and Swiss samples and monomorphic in the Spanish ones. Concerning species B, 20 mutations were polymorphic in French samples and monomorphic in CEE ones, but only nine showed the reverse pattern. In addition, the average number of nucleotide differences between populations was shown to be 0.330 between France and Spain that shared three mutations; 1.292 between France and Italy and Switzerland combined that shared one mutation; 1.401 between Spain and Italy and Switzerland combined that shared no mutation for species A; and 0.404 between France and CEE for species B that shared four mutations.

Finally, Table 2 shows that the value of Tajima’s D for the COI-t and ITS2 genes for species A is non-significant. For species B, the COI-t gene has a Tajima’s D value that is both negative and significant, indicating an excess of rare alleles. For ITS2, Tajima’s D is not significant.

### 3.3 Evolutionary history of the *C. pruni* complex

The ABC analyses provided a posterior probability for each hypothetical scenario and each replicate. Mean posterior probabilities of scenarios, 95% confidence intervals (CI), and the number of times the scenario was inferred as the best out of ten replicates are given in Table 3.

**Table 3:** Results of the ABC analyses. For each scenario (sc) or hypothesis, the mean posterior probability of this scenario based on a logistic regression, the mean confidence interval at 95% and the number of times it was inferred as the best scenario is given.

| Hypotheses | Mean posterior probability<br>of scenario (logistic<br>regression) | Mean confidence<br>interval (95%) | number of time inferred as<br>the best scenario |
| --- | --- | --- | --- |
| **Step A : Population Divergence** |  |  |  |
| Sc1 | 0.0232 | [0.009832 ; 0.04843] | 0/10 |
| Sc2 | 0.03332 | [0.01973 ; 0.05701] | 0/10 |
| **Sc3** | **0.4923** | **[0.47519 ; 0.50942]** | **9/10** |
| Sc4 | 0.23069 | [0.20461 ; 0.25677] | 1/10 |
| Sc5 | 0.22049 | [0.19901 ; 0.24197] | 0/10 |
| **Step B : Interspecific Admixture** |  |  |  |
| Sc1 | 0.42124 | [0.40333 ; 0.43915] | 0/10 |
| **Sc2** | **0.57876** | **[0.56085 ; 0.59667]** | **10/10** |
| **Step C : Intraspecific Admixture** |  |  |  |
| Sc1 | 0.23038 | [0.21559 ; 0.2452] | 2/10 |
| Sc2 | 0.25325 | [0.23751 ; 0.26899] | 2/10 |
| Sc3 | 0.25982 | [0.24185 ; 0.27777] | 3/10 |
| Sc4 | 0.25656 | [0.24015 ; 0.27294] | 3/10 |
| **Step D : Corsica Divergence** |  |  |  |
| Sc1 | 0.03463 | [0.02374 ; 0.0627] | 0/10 |
| **Sc2** | **0.47782** | **[0.44767 ; 0.51096]** | **6/10** |
| Sc3 | 0.04344 | [0.03043 ; 0.07274] | 0/10 |
| Sc4 | 0.04679 | [0.03523 ; 0.0755] | 1/10 |
| Sc5 | 0.33757 | [0.30853 ; 0.36662] | 3/10 |
| Sc6 | 0.05874 | [0.04328 ; 0.0877] | 0/10 |

The first ABC analysis (step A, Fig. 1) that tested five alternative divergence scenarios showed the highest support for scenario 3, which obtained the highest mean posterior probability (0.4923) with a mean 95% confidence interval ranging between 0.47519 and 0.50942, and was inferred as the best in nine out of ten cases. Scenario 3 assumes that the divergence between species A and B occurred in what is actually known as France, and that Spanish and CEE populations diverged independently from A and B French populations, respectively. All other models showed much lower posterior probabilities in comparison, especially scenarios 1 and 2, which could be ruled out as being incompatible with the data (Gilabert et al. 2023). Those scenarios assumed that the ancestral population was the Spanish one (sc1) and the Central and Eastern European one (sc2), respectively, instead of the French one, and that the divergence and expansion occurred in a west-to-east (sc1) or east-to-west (sc2) fashion. There was only one exception in which our tests inferred scenario 4 as the best (Table 3). Thereafter, scenario 3 appeared as the most robust hypothesis and was used as a backbone to design the following analysis and scenarios investigating admixture and divergence of the Corsican population.

In the second ABC analysis (step B, Fig. 1), two scenarios that assumed a presence and absence of admixture between species A and B, respectively, after their divergence were compared. All our replicates inferred scenario 2, assuming no admixture between both species after their divergence and thus identical to scenario 3 of step A, to be the most likely. It showed a mean posterior probability of 0.57876 and a 95% CI ranging from 0.56085 to 0.59667 (Table 3).

For the third ABC analysis (step C, Fig. 1), four competing scenarios were tested to determine if there had been admixture between the French and Spanish populations of species A, and between the French and CEE populations of species B. This step revealed no clear advantage of one scenario over another. Both scenario 3 and 4 were inferred as the best, three out of ten times (mean posterior probability = 0.25982 and 0.25656, respectively), and scenario 1 and 2 were the best choices, two out of ten times (mean posterior probability = 0.23038 and 0.25325, respectively) (Table 3). It is therefore impossible to conclude about the presence or absence of admixture between populations of the same species in the evolutionary history of these populations based on these data and analyses. For the sake of parsimony, we considered that there was no admixture in the following step.

Finally, in the last ABC analysis (step D, Fig. 1), six competing scenarios were considered to model the divergence of the Corsican population (species B) from the French or Italian ones. Four scenarios (sc1, 3, 4 & 6), all assuming divergence before/during separation of Corsica from the continent (15-28 Mya) and testing the presence or absence of admixture during the Messinian salinity crisis, and considering divergence from French or Italian populations, had very low posterior probabilities and were never inferred as the best scenario in our analyses (except one time for sc4). They can therefore be eliminated. Aside from that, the strongest support was obtained for scenario 2 in six out of ten replicates, with a mean posterior probability of 0.47782 (95% CI = 0.44767 to 0.51096). Moreover, scenario 5 was inferred as the best every other time except one (mean posterior probability = 0.33757) (Table 3). Both these scenarios inferred a divergence of the Corsican population during the Messinian salinity crisis (5.33-5.96 Mya). Scenario 2 inferred that the Corsican population diverged from other French populations, whereas scenario 5 implied that they diverged from CEE populations (most probably Italian ones in this case). We conclude that the most likely scenario seems to be a divergence of the Corsican population during the Messinian salinity crisis (5.33-5.96 Mya), probably from French populations with a non-excludable possibility that they diverged from CEE populations.

Because the analyses performed during the D step do not seem to be able to compute satisfactory posterior distributions of parameters, we only present those posterior distributions for scenario 3 of step A since it was considered to be the best one, and the selected scenario of steps B and C were identical to this scenario (Table 4).

**Table 4:** Mean parameter estimates for the best population divergence scenario (scenario 3, step A, Fig. 1) Values marked with an * were inferred from step D, scenario 2 (Fig. 1). For each parameter, its name, mean, median, 25% and 75% quartile values are given.

| Parameter | Parameter name | Mean | Median | q025 | q975 |
| --- | --- | --- | --- | --- | --- |
| Effective size of S population | NS | 3,64E+05 | 8,13E+05 | 8,96E+04 | 6,95E+05 |
| Effective size of FA population | NFA | 1,68E+05 | 7,92E+04 | 1,63E+03 | 5,61E+05 |
| Effective size of FB population | NFB | 2,16E+05 | 1,14E+05 | 2,43E+03 | 6,61E+05 |
| Effective size of CEE population | NCEE | 5,72E+06 | 2,47E+05 | 1,19E+04 | 6,50E+05 |
| Effective size of Corsica population | NC* | 5,51E+05 | 5,59E+05 | 4,70E+05 | 5,71E+05 |
| Divergence time between A & B species | tdiv | 2,09E+07 | 2,10E+07 | 1,62E+07 | 2,65E+07 |
| Divergence time of S population from FA | tS | 6,61E+06 | 5,11E+06 | 4,06E+06 | 1,03E+07 |
| Divergence time of CEE population from FB | tCEE | 6,36E+06 | 4,96E+06 | 4,04E+06 | 1,00E+07 |
| Divergence time of Corsica population | tc* | 5,69E+06 | 5,69E+06 | 5,66E+06 | 5,69E+06 |
| Duration of the bottleneck | db | 2,87E+03 | 2,18E+03 | 1,12E+03 | 8,72E+03 |
| Effective size of S population during the bottleneck | NSb | 1,81E+02 | 8,92E+01 | 1,13E+01 | 7,56E+02 |
| Effective size of FA population during the bottleneck | NFAb | 2,64E+02 | 1,83E+02 | 1,21E+02 | 8,28E+02 |
| Effective size of FB population during the bottleneck | NFBb | 2,57E+02 | 1,72E+02 | 1,21E+02 | 8,10E+02 |
| Effective size of CEE population during the bottleneck | NCEEb | 2,02E+02 | 1,19E+02 | 1,19E+01 | 7,72E+02 |
| Effective size of Corsica population during the bottleneck | NCb* | 4,73E+02 | 4,56E+02 | 4,35E+02 | 5,67E+02 |
| Effective size of the origin population | NO | 1,89E+05 | 1,19E+05 | 1,11E+05 | 7,04E+05 |

## 4 – Discussion

The main objective of this study was to decipher the evolutionary history of the two cryptic species of the complex *C. pruni* in Europe using phylogenetic and population genetic methods. This allowed us to identify the most probable source population and reconstruct the migratory routes of these insects, vectors of a phytoplasma causing the ESFY on *Prunus* trees.

### 4.1 Evolutionary history of the *C. pruni* complex within the Western Palearctic

To our knowledge, this study is the first attempt at reconstructing the evolutionary history of a psyllid at a continental scale. Previous studies have been conducted on an island scale (Canary Island archipelago: Percy 2003a; Hawaii: Hembry et al. 2021). Before the present analyses, it could have been expected that the formation of this cryptic pair of species would be a classic case of glacial refugia in the Iberian Peninsula, followed by a recolonisation of Europe from west to east, as recurrently identified in Europe (Taberlet et al. 1998; Hewitt 2000; Hewitt 2001; Hewitt 2004). Surprisingly, the present ABC analysis showed strong posterior support for the scenario where the two cryptic species diverged in France, their origin population, from a common ancestor, approximately 20.19 Mya (95% CI: 24.26-28.59 Mya), before expanding into Spain and Central and Eastern Europe around 6.61 and 6.36 Mya (95% CI: 4.04-10.3 Mya), respectively, with species B passing into Corsica during the Messinian salinity crisis (5.33-5.96 Mya; Krijgsman et al. 1999) from either French or Italian populations (Fig. 6). We also found no admixture between the two species after divergence, which is in agreement with recent studies that showed no trace of admixture between both species of the complex (Peccoud et al. 2018; Sauvion et al. 2024), but could not conclude as to the presence or absence of admixture between populations of the same species. Thus, the ABC method helped to clarify the Western Palearctic history of *C. pruni*, even though some of the results still need to be considered with caution.

**Fig. 6:**
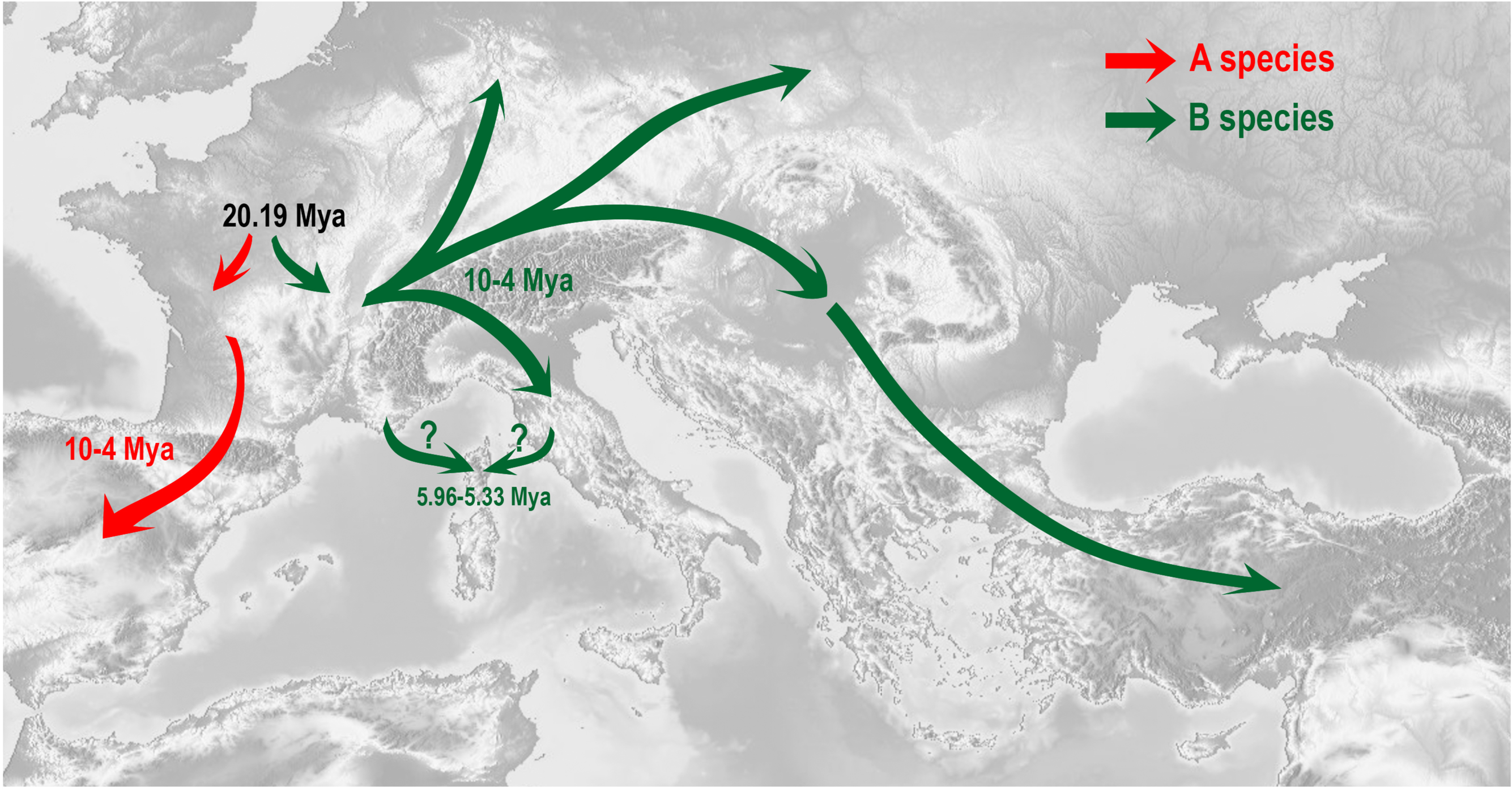
Summary map of the best scenario inferred with the ABC analyses of the evolutionary history of *C. pruni*.

### 4.2 ABC results are congruent with the actual genetic diversity and structure of psyllids

The results of the ABC analysis are very congruent with the results from the preliminary population genetic analyses. For example, the characterisation of the genetic diversity of the *C. pruni* complex in Europe using the COI-t and ITS2 genes has shown that French populations presented more mutations and higher haplotype and nucleotide diversity than Spanish ones for species A, followed by CEE populations for species B. In addition, French populations showed a higher number of private haplotypes than other geographic areas, even when taking account of the unequal sample sizes between samples with a rarefaction approach. This was also confirmed by the visual analysis of the haplotype networks and distribution maps for both genes and both species. Moreover, the French populations had the most mutations that were polymorphic within themselves but monomorphic within other populations. All these results point toward France being the origin population for the two species, and the Spanish and CEE populations that evolved from French populations of species A and B, respectively.

Furthermore, the analyses showed that the values of Tajima’s D for the COI-t gene in species B were negative and significant for both French and Central and Eastern European populations, as well as for the French populations for species A, whereas they were non-significant for Spanish and Italian/Swiss populations of species A. This suggests that while species A evolved in a neutral manner (without selective pressure) – except maybe from the French populations – the B species showed an excess of rare alleles that could indicate a fast expansion after a bottleneck, which would be consistent with a scenario of fast expansion of the CEE population from the French ones. In this case, the excess of rare alleles observed in both species in the French populations could be the result of (i) a population expansion after their divergence, or (ii) a recent selective sweep or the linkage of the COI-t gene to a swept gene. Furthermore, seeing that Tajima’s D values are positive and non-significant for the ITS2 gene, for which we only sampled French populations, we could argue that the excess of rare alleles observed for the COI-t gene in the French populations is probably more of an artefact due to the linkage to a swept gene or a selective sweep of the COI-t gene than the result of an expansion, which would probably have resulted in Tajima’s D being negative for the ITS2 gene as well.

### 4.3 ABC analyses: plausibility and limits

In the ABC analysis, some of the scenario choices were associated with low uncertainty. For instance, for steps A and B, the best scenario was inferred in 9/10 and 10/10 replicates with consistently high posterior probabilities. Alternate scenarios were associated with such low posterior probabilities that they could be refuted (as advocated in Gilabert et al. 2023). This suggests that the variability revealed at marker COI-t on the studied populations is informative enough to allow us to discriminate between the different scenarios based on the summary statistics used. In contrast, some of the results should be considered with more caution. In particular, the scenario choice in steps C and D led to moderate posterior probabilities and inconsistencies in scenario ranking across replicates. Similarly, posterior estimates of the parameters in step D scenarios could not be obtained (due to large confidence intervals and posterior distributions very close to prior distributions). These limitations probably lie in the use of a single marker gene for the analysis, as well as the limited number of sites sampled outside of France. Indeed, the key to a successful application of ABC methods resides in the capacity of the summary statistics to encompass the patterns and properties of the data. Thus, even if ABC analyses have the potential to use very complex models, if this complexity cannot be supported by the data, the analysis will not provide additional information (Nielsen & Beaumont 2009). Furthermore, the Bayesian estimation of parameters may be improved by the combination of multiple datasets with several types of markers (Wegmann & Excoffier 2010). Thus, even if our analyses made it possible to decipher the evolutionary history of the *C. pruni* complex in the Western Palearctic, the present dataset could definitely be improved by genotyping other genes (e.g., microsatellites, other nuclear or cytoplasmic genes), and obtaining a better representation of geographical areas that could not be sampled for this study (e.g., sample sites outside of France for the ITS2 gene).

### 4.4 Dating the psyllid tree with host phylogeny

In the absence of sufficient and informative fossil data, we have attempted to calibrate the Psyllinae phylogenic tree on the basis of molecular dating of host plants (Rosaceae). A criticism of this approach might be that the present phylogenetic tree has been built from a single gene. However, the resulting phylogeny is highly congruent with the current consensual hypothesis on the phylogeny of Psylloidea based on complete mitogenome or multilocus DNA sequences (Percy et al. 2018; Cho et al. 2019; Burckhardt et al. 2021). Moreover, the literature survey of dated phylogenies of Rosaceae used to calibrate our nodes showed very congruent timetrees across studies. Thus, the only grey area remaining is the accuracy of the underlying hypothesis used to calibrate some nodes of the tree. It can be inferred that, at least in the Psyllinae family, there were co-evolution or co-speciation events between psyllids and their hosts, or at least that the speciation of a host plant was quickly followed by the speciation of its specialised psyllid (Hodkinson 1984; White & Hodkinson 1985; Brown & Hodkinson 1988; Van Klinken 2000; Percy et al. 2004). Yet, we cannot exclude the hypothesis of a host switch that would not have been contemporaneous to host divergence (Percy et al. 2004; Ouvrard et al. 2015). In that case, the divergence date between species A and B used in the ABC analysis could have been incorrect. However, preliminary tests (not shown) revealed that, apart from parameter values, the results of scenario choice analysis (origin population, order of divergence, which population diverged from which, presence or absence of admixture, etc.) did not depend to any great degree on the prior divergence date used. To confirm these results, it would be interesting to have access to other phylogenetic studies on the Psyllinae based on other methods (e.g., newly discovered fossils or molecular clock-based analyses).

### 4.5 Biological factors explaining past divergence and expansion

The fact that the divergence date between both cryptic species found in this study is ancient is consistent with the strong reproductive barriers and almost complete reproductive isolation previously shown by Peccoud et al. (2018). This could thus be a case of cryptic species in stasis (in opposition to a recent divergence of an evolutionary convergence), as described by Struck et al. (2018), also referred to as the “ecological niche conservatism hypothesis” or morphological stase by Fišer et al. (2018). Under this hypothesis, cryptic species maintain a high degree of morphological similarity over a long period of time, which could be the result of a lack of genetic variation, a strong stabilising selection, very strong gene flux overwhelming the local adaptations or pleiotropic effects constraining the adaptation, or a combination of several of these factors (Struck et al. 2018; Fišer et al. 2018). Several cryptic species showing this phenomenon have already been described, e.g., in *Niphargus fongi* (Fišer & Zagmajster, 2009) and in two Leucothoid amphipod species (Richards et al. 2012).

As previously mentioned, the most likely hypothesis to explain why the species diverged would be a host switch, given that host switching to a closely related host is one of the most common conditions associated with psyllid speciation (Percy 2003a; Percy et al. 2004; Ouvrard et al. 2015). In this case, both species of *C. pruni* can be found in strict sympatry on *Prunus* trees (mainly blackthorn, *Prunus spinosa* L., and plum, *Prunus domestica* L.), which may cast doubts on the host switch hypothesis. However, it is commonly admitted that *Prunus* trees are not the only host plant of *C. pruni* during its life cycle. Indeed, *C. pruni* females reproduce and lay eggs on *Prunus* trees in the spring and then migrate to sites where they overwinter on coniferous trees (formally shelter plants; Thébaud et al. 2009). A plausible hypothesis for the divergence of both *C. pruni* species could therefore be a host switch, not of their *Prunus* host plant, but of their coniferous shelter plant. This hypothesis is also supported by the observation that species A prefers pine, while species B prefers spruce or fir (Sauvion et al. 2024 in prepar).

After divergence, the expansion of the *C. pruni* complex seems to have expanded across Europe. A hypothesis to explain this change in the species distribution could be a climate-induced host expansion or a change in climate suitability in Spain and CEE. The divergence time between French, Spanish and CEE populations was dated at around 6.61 and 6.36 Mya, respectively, thus during the late Neogene, in the Upper Miocene (11.61 to 5.33 Ma) or, more precisely, the Messinian period (around 7.3-5.2 Mya). During this period, it was shown that the Iberian Peninsula went through significant changes in both vegetation and sea level, probably triggered by orbital-scale climate change. The climate was warm and dry in this region, whereas in the southwest of France, the Black Sea and the north of Italy, it was warm and humid (Jimenez-Moreno et al. 2013; Prista et al. 2015). In addition, a progressive reduction in the most hermophilous and high-water requirement plants and an increase in seasonal-adapted plants from higher altitude belts, including mesothermic elements, altitudinal trees and herbs, appears to have occurred in Southeast Europe during this period (Jimenez-Moreno et al. 2007).

## Author contributions

NS conceptualised the project with support from VR. NS provided all the psyllid specimens and related sequences used for phylogenetic analyses. MD performed all analyses and simulations. NS and VR approved the results. MD and NS wrote the initial version and all authors contributed to revising subsequent versions. All authors approved of the content of the final manuscript.

## Acknowledgements

The authors acknowledge the support of funding from Labex Agro – Montpellier project E-SPACE (contract # 1504-004), and the French National Research Agency (ANR) for the BEYOND project (contract # 20-PCPA-0002), which facilitated the regular interactions of all authors and led to the conception and realisation of this publication. We are very grateful to Josiane Peyre for her valuable technical assistance. A large part of the molecular sequencing for this study was supported by the SPEED@ID (Accurate SPEciEs Delimitation and IDentification of Eukaryotic biodiversity using DNA markers) project funded by F-BoL, the French Barcode of Life initative - Genoscope Evry-France; and PHYLOPSYL from the project “Bibliothèque du vivant” (BdV) funded by three French institutions (the CNRS, INRAE and MNHN).

## Data availability statement

Fig. S1: Maps of sampled sites

Each site number corresponds to the site number in Table S1. Sample sites with only the A species present are represented in red, the ones with only the B species present are represented in green, and the ones with both species present in sympatry are represented in black.

Fig. S2: Maximum likelihood phylogenetic tree based on the COI-tRNA-COII mitochondrial gene of 94 Psyllinae species with annotations of the molecular dating analysis.

Black circles represent the seven nodes that were calibrated for the analysis; names in light green represent the host plant of the species next to it; names in brown represent the geographic localisation of the host plant; other colours are associated with clades. Calibration dates for the seven calibrated nodes and associated references are the following:

| Node | « Uniform »<br>distribution | « Normal »<br>distribution = mean ;<br>standard dev | References : |
| --- | --- | --- | --- |
| 1 | 77.3 – 91.02 Ma | 87 Ma ; 5 Ma | Zhang (2017); Chin (2014) |
| 2 | 30 – 72.1 Ma | 50 Ma ; 10 Ma | Zhang (2017); Chin (2014); Du (2021) |
| 3 | 30 – 72.1 Ma | 50 Ma ; 10 Ma | Zhang (2017); Chin (2014); Du (2021) |
| 4 | 25.23 – 33.35 Ma | 27.49 Ma ; 3 Ma | Korotkova (2018) |
| 5 | 45.63 – 92.84 Ma | 70 Ma ; 12 Ma | Zhang (2017); Chin (2014); Du (2021) |
| 6 | 16 – 350 Ma | 16 – 350 Ma | Klimaszewski (1993); Becker (1964); Johnson (2018) |
| 7 | 1 – 16 Ma | 1 – 16 Ma | Percy (2003a); Percy (2003b) |

Fig. S3: Maximum likelihood phylogenetic tree based on the COI-tRNA-COII mitochondrial gene of 94 Psyllinae species obtained with MEGA.

Fig. S4: FastME (distance-based) phylogenetic tree based on the COI-tRNA-COII mitochondrial gene of 94 Psyllinae species obtained with NGphylogeny.

Fig. S5: Raw and complete dated phylogeny (ML) of the Psyllinae species based on the COI-tRNA-COII mitochondrial gene of 94 Psyllinae species obtained with MEGA.

External group not shown.

Table S1: List and description of sample sites.

For each sample site, the number of the site, its locality, the species present, its latitude and longitude as well as the number of sequences for the COI-t and ITS2 genes are given.

Table S2: Names of the species used for phylogenetic analysis and their related NCBI-accession for the cox1-trnL-cox2 gene.

Table S3: Maximum Likelihood fits of 24 different nucleotide substitution models in MEGA.

For each model, BIC (Bayesian Information Criterion), AICc value (Akaike Information Criterion, corrected), Maximum Likelihood value (lnL), and the number of parameters (including branch lengths) are presented. Whenever applicable, estimates of gamma shape parameters and/or the estimated fraction of invariant sites are shown. Assumed or estimated values of transition/transversion bias (R) are shown for each model as well. They are followed by nucleotide frequencies (f) and rates of base substitutions (r) for each nucleotide pair. Relative values of instantaneous r should be considered when evaluating them. For simplicity, the sum of r values is set equal to 1 for each model. For estimating ML values, a tree topology was automatically computed. This analysis involved 94 nucleotide sequences.

Table S4: Summary statistics used in the ABC analyses.

