## Supplemental Materials Figs+Table for "Origin, divergence and migration routes of psyllids of the *Cacopsylla pruni* complex (Hemiptera: Psylloidea) inferred by Approximate Bayesian Computation methods"

Figure S1

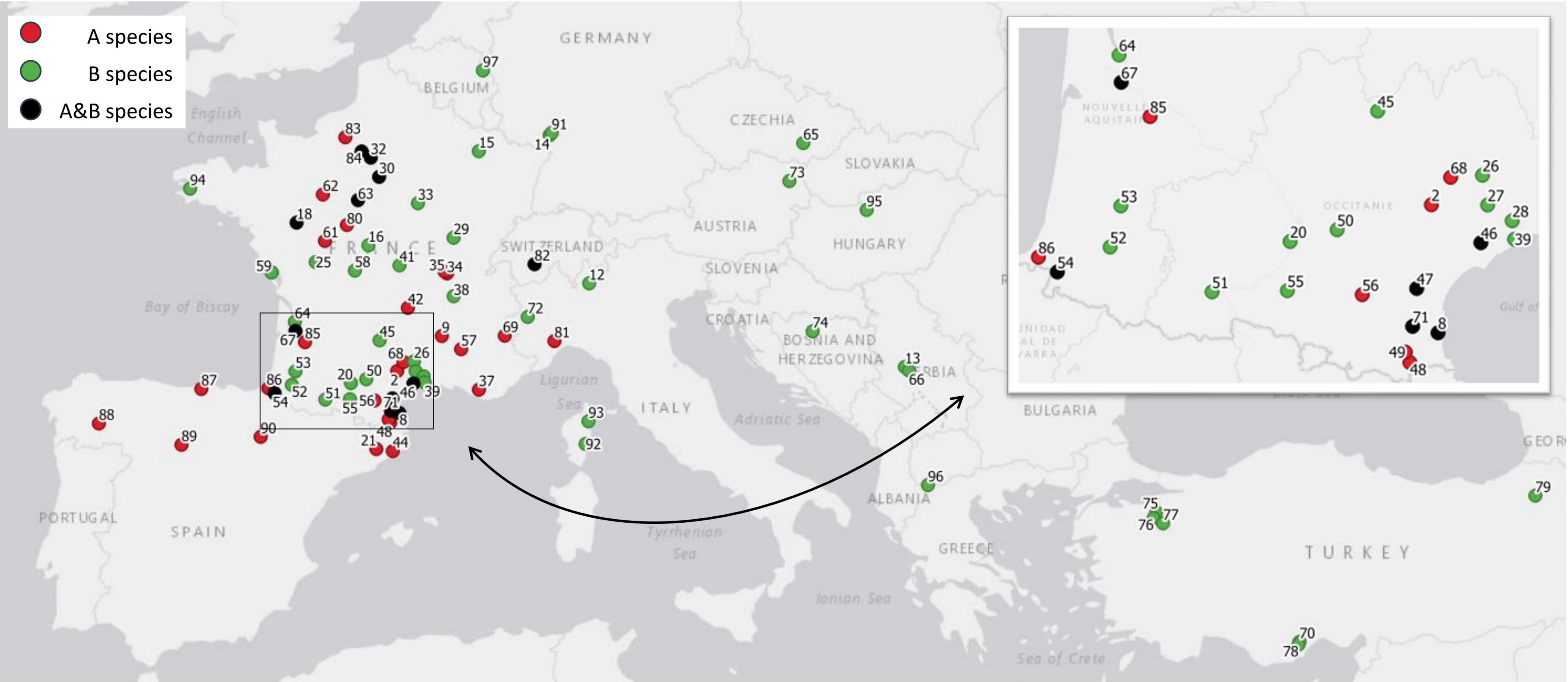

A species
B species
A&B species
GERMANY
BELGIUM
CZECHIA
SLOVAKIA
AUSTRIA
HUNGARY
SWITZERLAND
SLOVENIA
CROATIA
BOSNIA AND HERZEGOVINA
SERBIA
BULGARIA
ALBANIA
GREECE
TURKEY
ITALY
SPAIN
PORTUGAL
FRANCE
English Channel
Bay of Biscay
Ligurian Sea
Adriatic Sea
Tyrrhenian Sea
Ionian Sea
Sea of Crete
NOUVELLE AQUITAINE
OCCITANIE


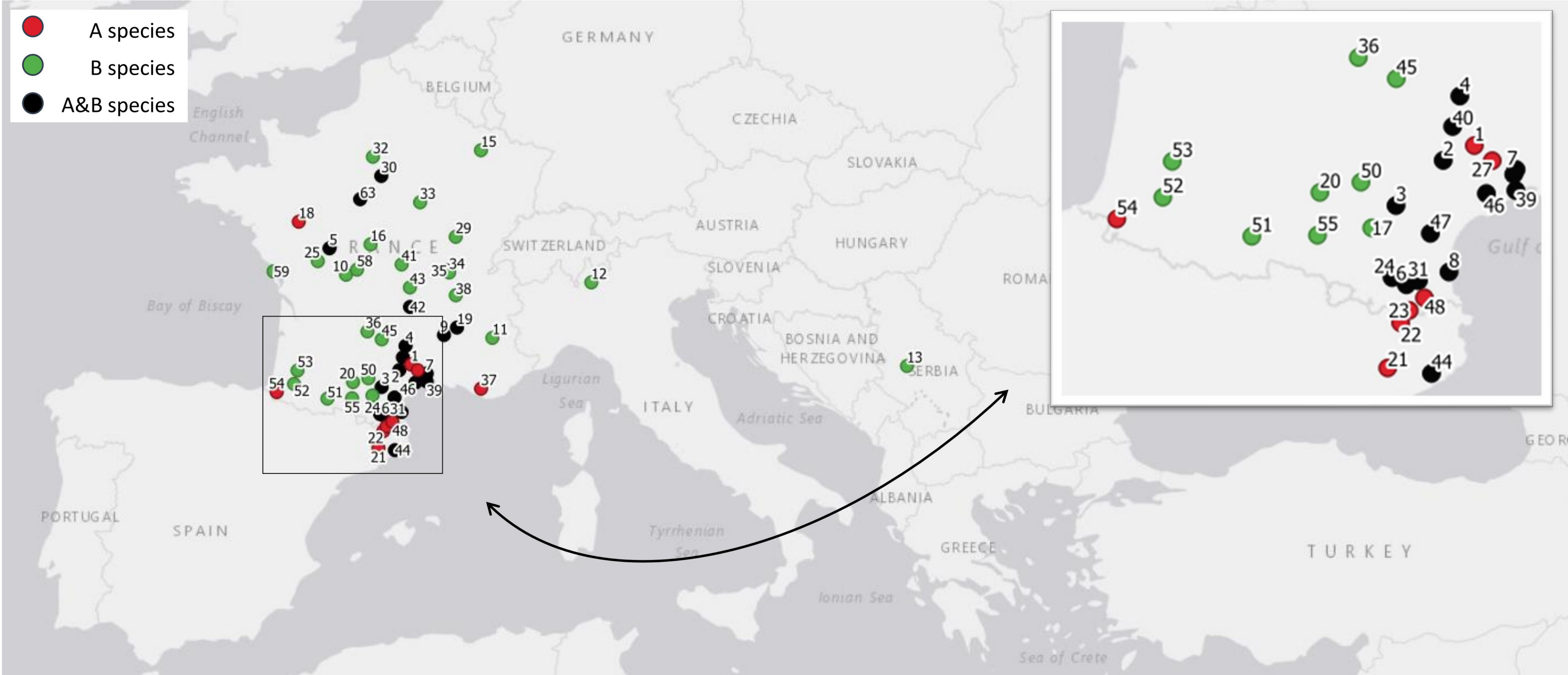

A species
B species
A&B species
GERMANY
BELGIUM
CZECHIA
SLOVAKIA
AUSTRIA
HUNGARY
SWITZERLAND
SLOVENIA
CROATIA
BOSNIA AND HERZEGOVINA
SERBIA
ALBANIA
GREECE
TURKEY
ITALY
SPAIN
PORTUGAL
English Channel
Bay of Biscay
Ligurian Sea
Adriatic Sea
Tyrrhenian Sea
Ionian Sea
Sea of Crete

Figure S2

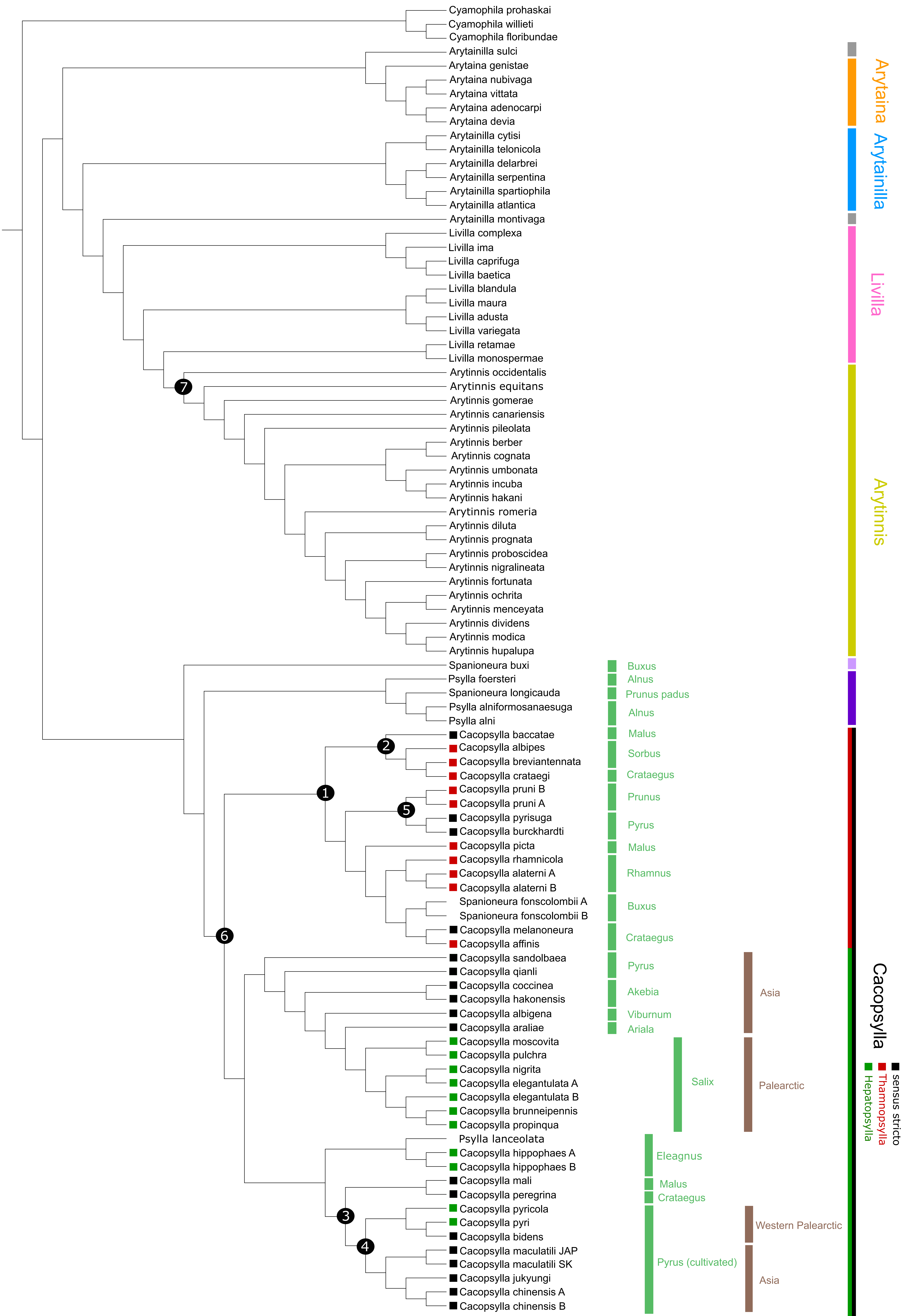

Cyamophila prohaskai
Cyamophila willieti
Cyamophila floribundae
Arytainilla sulci
Arytaina genistae
Arytaina nubivaga
Arytaina vittata
Arytaina adenocarpi
Arytaina devia
Arytainilla cytisi
Arytainilla telonicola
Arytainilla delarbrei
Arytainilla serpentina
Arytainilla spartiophila
Arytainilla atlantica
Arytainilla montivaga
Livilla complexa
Livilla ima
Livilla caprifuga
Livilla baetica
Livilla blandula
Livilla maura
Livilla adusta
Livilla variegata
Livilla retamae
Livilla monospermae
Arytinnis occidentalis
Arytinnis equitans
Arytinnis gomerae
Arytinnis canariensis
Arytinnis pileolata
Arytinnis berber
Arytinnis cognata
Arytinnis umbonata
Arytinnis incuba
Arytinnis hakani
Arytinnis romeria
Arytinnis diluta
Arytinnis prognata
Arytinnis proboscidea
Arytinnis nigralineata
Arytinnis fortunata
Arytinnis ochrita
Arytinnis menceyata
Arytinnis dividens
Arytinnis modica
Arytinnis hupalupa
Spanioneura buxi
Psylla foersteri
Spanioneura longicauda
Psylla alniformosanaesuga
Psylla alni
Cacopsylla baccatae
Cacopsylla albipes
Cacopsylla breviantennata
Cacopsylla crataegi
Cacopsylla pruni B
Cacopsylla pruni A
Cacopsylla pyrisuga
Cacopsylla burckhardti
Cacopsylla picta
Cacopsylla rhamnicola
Cacopsylla alaterni A
Cacopsylla alaterni B
Spanioneura fonscolombii A
Spanioneura fonscolombii B
Cacopsylla melanoneura
Cacopsylla affinis
Cacopsylla sandolbaea
Cacopsylla qianli
Cacopsylla coccinea
Cacopsylla hakonensis
Cacopsylla albigena
Cacopsylla araliae
Cacopsylla moscovita
Cacopsylla pulchra
Cacopsylla nigrita
Cacopsylla elegantulata A
Cacopsylla elegantulata B
Cacopsylla brunneipennis
Cacopsylla propinqua
Psylla lanceolata
Cacopsylla hippophaes A
Cacopsylla hippophaes B
Cacopsylla mali
Cacopsylla peregrina
Cacopsylla pyricola
Cacopsylla pyri
Cacopsylla bidens
Cacopsylla maculatili JAP
Cacopsylla maculatili SK
Cacopsylla jukyungi
Cacopsylla chinensis A
Cacopsylla chinensis B
1
2
3
4
5
6
7
Buxus
Alnus
Prunus padus
Alnus
Malus
Sorbus
Crataegus
Prunus
Pyrus
Malus
Rhamnus
Buxus
Crataegus
Pyrus
Akebia
Viburnum
Ariala
Salix
Eleagnus
Malus
Crataegus
Pyrus (cultivated)
Asia
Palearctic
Western Palearctic
Asia
Arytaina
Arytainilla
Livilla
Arytinnis
Cacopsylla
sensus stricto
Thamnopsylla
Hepatopsylla

Figure S3

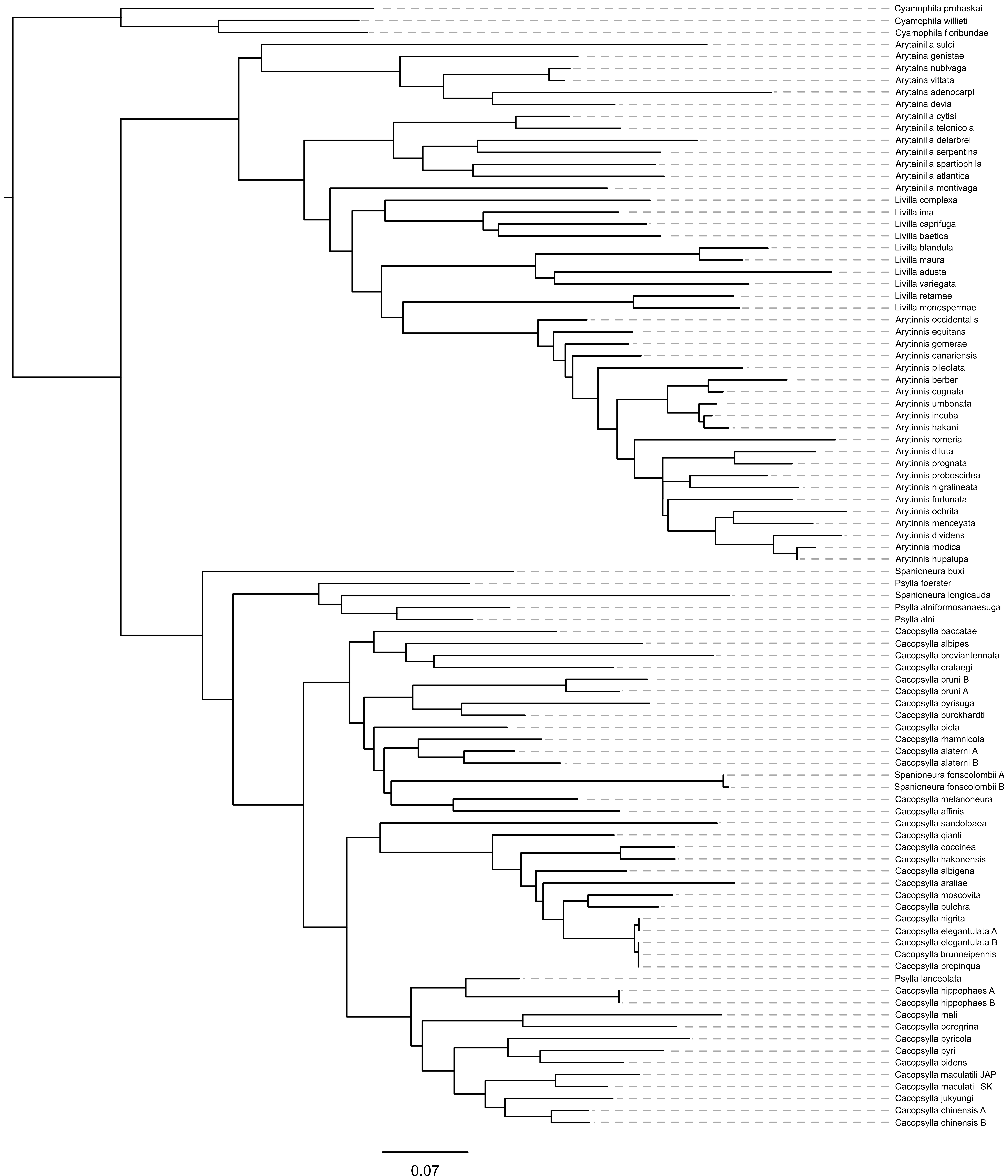
Cyamophila prohaskai
Cyamophila willieti
Cyamophila floribundae
Arytainilla sulci
Arytaina genistae
Arytaina nubivaga
Arytaina vittata
Arytaina adenocarpi
Arytaina devia
Arytainilla cytisi
Arytainilla telonicola
Arytainilla delarbrei
Arytainilla serpentina
Arytainilla spartiophila
Arytainilla atlantica
Arytainilla montivaga
Livilla complexa
Livilla ima
Livilla caprifuga
Livilla baetica
Livilla blandula
Livilla maura
Livilla adusta
Livilla variegata
Livilla retamae
Livilla monospermae
Arytinnis occidentalis
Arytinnis equitans
Arytinnis gomerae
Arytinnis canariensis
Arytinnis pileolata
Arytinnis berber
Arytinnis cognata
Arytinnis umbonata
Arytinnis incuba
Arytinnis hakani
Arytinnis romeria
Arytinnis diluta
Arytinnis prognata
Arytinnis proboscidea
Arytinnis nigralineata
Arytinnis fortunata
Arytinnis ochrita
Arytinnis menceyata
Arytinnis dividens
Arytinnis modica
Arytinnis hupalupa
Spanioneura buxi
Psylla foersteri
Spanioneura longicauda
Psylla alniformosanaesuga
Psylla alni
Cacopsylla baccatae
Cacopsylla albipes
Cacopsylla breviantennata
Cacopsylla crataegi
Cacopsylla pruni B
Cacopsylla pruni A
Cacopsylla pyrisuga
Cacopsylla burckhardti
Cacopsylla picta
Cacopsylla rhamnicola
Cacopsylla alaterni A
Cacopsylla alaterni B
Spanioneura fonscolombii A
Spanioneura fonscolombii B
Cacopsylla melanoneura
Cacopsylla affinis
Cacopsylla sandolbaea
Cacopsylla qianli
Cacopsylla coccinea
Cacopsylla hakonensis
Cacopsylla albigena
Cacopsylla araliae
Cacopsylla moscovita
Cacopsylla pulchra
Cacopsylla nigrita
Cacopsylla elegantulata A
Cacopsylla elegantulata B
Cacopsylla brunneipennis
Cacopsylla propinqua
Psylla lanceolata
Cacopsylla hippophaes A
Cacopsylla hippophaes B
Cacopsylla mali
Cacopsylla peregrina
Cacopsylla pyricola
Cacopsylla pyri
Cacopsylla bidens
Cacopsylla maculatili JAP
Cacopsylla maculatili SK
Cacopsylla jukyungi
Cacopsylla chinensis A
Cacopsylla chinensis B
0.07

Figure S4

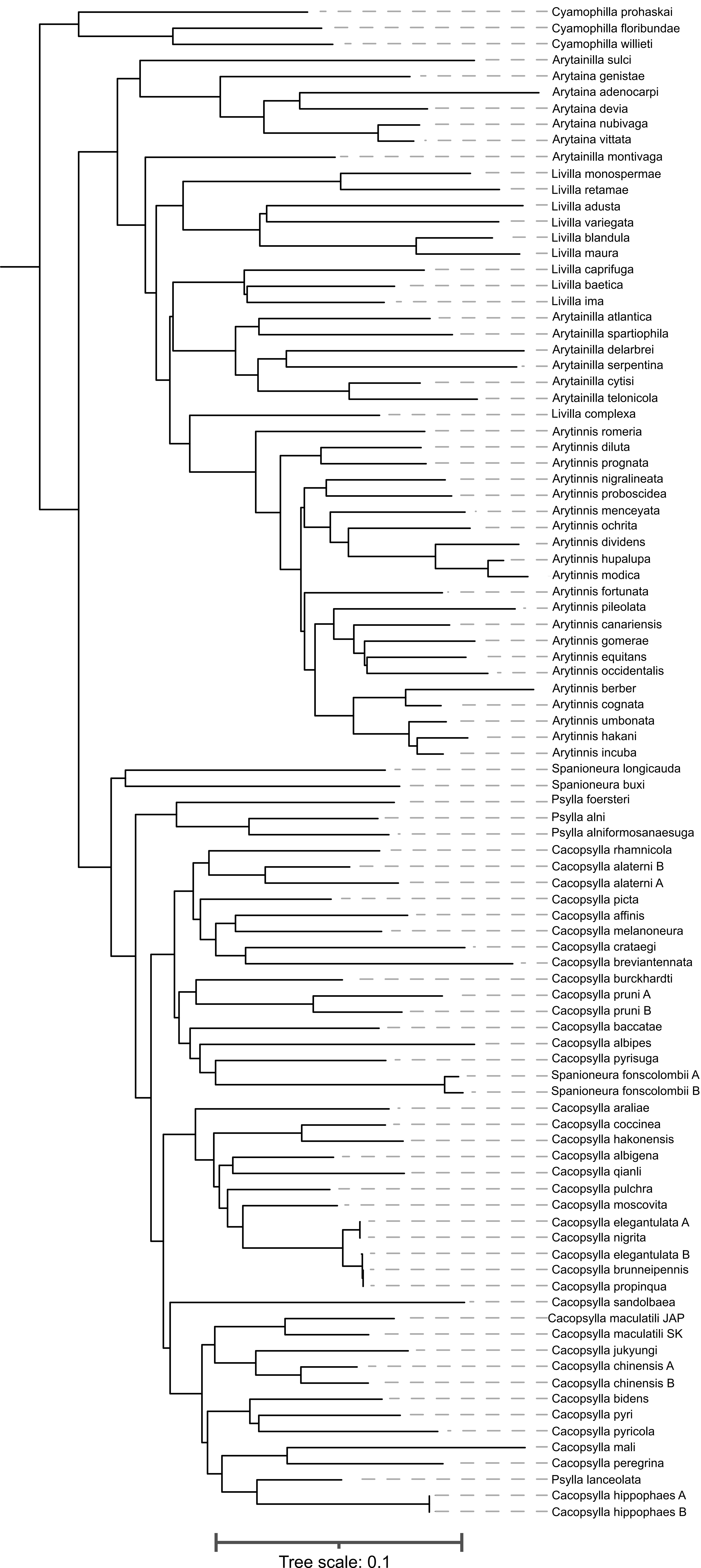

Cyamophilla prohaskai
Cyamophilla floribundae
Cyamophilla willieti
Arytainilla sulci
Arytaina genistae
Arytaina adenocarpi
Arytaina devia
Arytaina nubivaga
Arytaina vittata
Arytainilla montivaga
Livilla monospermae
Livilla retamae
Livilla adusta
Livilla variegata
Livilla blandula
Livilla maura
Livilla caprifuga
Livilla baetica
Livilla ima
Arytainilla atlantica
Arytainilla spartiophila
Arytainilla delarbrei
Arytainilla serpentina
Arytainilla cytisi
Arytainilla telonicola
Livilla complexa
Arytinnis romeria
Arytinnis diluta
Arytinnis prognata
Arytinnis nigralineata
Arytinnis proboscidea
Arytinnis menceyata
Arytinnis ochrita
Arytinnis dividens
Arytinnis hupalupa
Arytinnis modica
Arytinnis fortunata
Arytinnis pileolata
Arytinnis canariensis
Arytinnis gomerae
Arytinnis equitans
Arytinnis occidentalis
Arytinnis berber
Arytinnis cognata
Arytinnis umbonata
Arytinnis hakani
Arytinnis incuba
Spanioneura longicauda
Spanioneura buxi
Psylla foersteri
Psylla alni
Psylla alniformosanaesuga
Cacopsylla rhamnicola
Cacopsylla alaterni B
Cacopsylla alaterni A
Cacopsylla picta
Cacopsylla affinis
Cacopsylla melanoneura
Cacopsylla crataegi
Cacopsylla breviantennata
Cacopsylla burckhardti
Cacopsylla pruni A
Cacopsylla pruni B
Cacopsylla baccatae
Cacopsylla albipes
Cacopsylla pyrisuga
Spanioneura fonscolombii A
Spanioneura fonscolombii B
Cacopsylla araliae
Cacopsylla coccinea
Cacopsylla hakonensis
Cacopsylla albigena
Cacopsylla qianli
Cacopsylla pulchra
Cacopsylla moscovita
Cacopsylla elegantulata A
Cacopsylla nigrita
Cacopsylla elegantulata B
Cacopsylla brunneipennis
Cacopsylla propinqua
Cacopsylla sandolbaea
Cacopsylla maculatili JAP
Cacopsylla maculatili SK
Cacopsylla jukyungi
Cacopsylla chinensis A
Cacopsylla chinensis B
Cacopsylla bidens
Cacopsylla pyri
Cacopsylla pyricola
Cacopsylla mali
Cacopsylla peregrina
Psylla lanceolata
Cacopsylla hippophaes A
Cacopsylla hippophaes B
Tree scale: 0.1

Figure S5

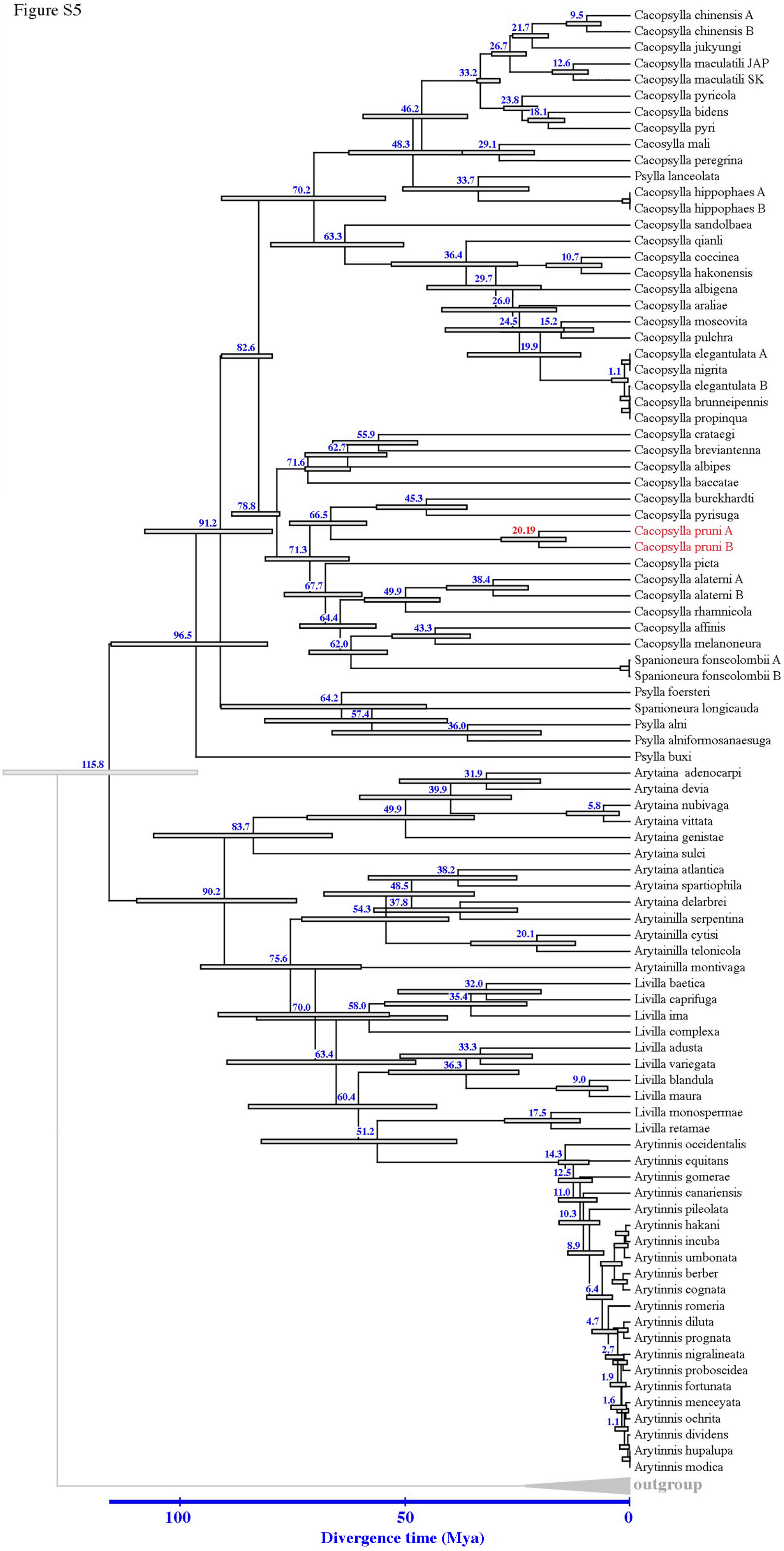
Cacopsylla chinensis A
Cacopsylla chinensis B
Cacopsylla jukyungi
Cacopsylla maculatili JAP
Cacopsylla maculatili SK
Cacopsylla pyricola
Cacopsylla bidens
Cacopsylla pyri
Cacosylla mali
Cacopsylla peregrina
Psylla lanceolata
Cacopsylla hippophaes A
Cacopsylla hippophaes B
Cacopsylla sandolbaea
Cacopsylla qianli
Cacopsylla coccinea
Cacopsylla hakonensis
Cacopsylla albigena
Cacopsylla araliae
Cacopsylla moscovita
Cacopsylla pulchra
Cacopsylla elegantulata A
Cacopsylla nigrita
Cacopsylla elegantulata B
Cacopsylla brunneipennis
Cacopsylla propinqua
Cacopsylla crataegi
Cacopsylla breviantenna
Cacopsylla albipes
Cacopsylla baccatae
Cacopsylla burckhardti
Cacopsylla pyrisuga
Cacopsylla pruni A
Cacopsylla pruni B
Cacopsylla picta
Cacopsylla alaterni A
Cacopsylla alaterni B
Cacopsylla rhamnicola
Cacopsylla affinis
Cacopsylla melanoneura
Spanioneura fonscolombii A
Spanioneura fonscolombii B
Psylla foersteri
Spanioneura longicauda
Psylla alni
Psylla alniformosanaesuga
Psylla buxi
Arytaina adenocarpi
Arytaina devia
Arytaina nubivaga
Arytaina vittata
Arytaina genistae
Arytaina sulci
Arytaina atlantica
Arytaina spartiophila
Arytaina delarbrei
Arytainilla serpentina
Arytainilla cytisi
Arytainilla telonicola
Arytainilla montivaga
Livilla baetica
Livilla caprifuga
Livilla ima
Livilla complexa
Livilla adusta
Livilla variegata
Livilla blandula
Livilla maura
Livilla monospermae
Livilla retamae
Arytinnis occidentalis
Arytinnis equitans
Arytinnis gomerae
Arytinnis canariensis
Arytinnis pileolata
Arytinnis hakani
Arytinnis incuba
Arytinnis umbonata
Arytinnis berber
Arytinnis cognata
Arytinnis romeria
Arytinnis diluta
Arytinnis prognata
Arytinnis nigralineata
Arytinnis proboscidea
Arytinnis fortunata
Arytinnis menceyata
Arytinnis ochrita
Arytinnis dividens
Arytinnis hupalupa
Arytinnis modica
outgroup
9.5
21.7
26.7
12.6
33.2
23.8
18.1
46.2
48.3
29.1
33.7
70.2
63.3
36.4
10.7
29.7
26.0
24.5
15.2
19.9
82.6
1.1
55.9
62.7
71.6
45.3
78.8
66.5
20.19
91.2
71.3
38.4
67.7
49.9
64.4
43.3
62.0
96.5
64.2
57.4
36.0
115.8
31.9
39.9
49.9
5.8
83.7
38.2
48.5
37.8
90.2
54.3
20.1
75.6
32.0
35.4
70.0
58.0
33.3
63.4
36.3
9.0
60.4
17.5
51.2
14.3
12.5
11.0
10.3
8.9
6.4
4.7
2.7
1.9
1.6
1.1
100
50
0
Divergence time (Mya)

Table S1

| Population number | Locality | Species | Latitude | Longitude | Number of ITS2 sequences | Number of COI-t sequences |
|---|---|---|---|---|---|---|
| 1 | Larzac | A | 43.9279 | 3.3117 | 37 | 0 |
| 2 | Coursegoules | A | 43.79 | 2.91 | 17 | 11 |
| 2 | Coursegoules | B | 43.79 | 2.91 | 7 | 0 |
| 3 | Haut-Languedoc | A | 3.3679 | 2.3016 | 8 | 0 |
| 3 | Haut-Languedoc | B | 3.3679 | 2.3016 | 14 | 0 |
| 4 | La Tieule | A | 44.387 | 3.1243 | 23 | 0 |
| 4 | La Tieule | B | 44.387 | 3.1243 | 4 | 0 |
| 5 | Montgamé | A | 46.7338 | 0.5102 | 1 | 0 |
| 5 | Montgamé | B | 46.7338 | 0.5102 | 25 | 0 |
| 6 | Prades | A | 42.6202 | 2.4356 | 13 | 0 |
| 6 | Prades | B | 42.6202 | 2.4356 | 2 | 0 |
| 7 | Prades-le-Lez | A | 3.7049 | 3.8502 | 30 | 0 |
| 7 | Prades-le-Lez | B | 3.7049 | 3.8502 | 4 | 0 |
| 8 | Torreilles | A | 42.7414 | 2.985 | 13 | 7 |
| 8 | Torreilles | B | 42.7414 | 2.985 | 14 | 1 |
| 9 | Fesche | A | 44.6528 | 4.4443 | 9 | 5 |
| 9 | Fesche | B | 44.6528 | 4.4443 | 3 | 0 |
| 10 | Bellac | B | 46.1068 | 1.0794 | 3 | 0 |
| 11 | Romette | B | 44.581994 | 6.106769 | 1 | 0 |
| 12 | Udine | B | 45.9235 | 9.5054 | 20 | 10 |
| 13 | Cacak | B | 43.8915 | 20.3502 | 9 | 3 |
| 14 | Neustadt | B | 49.3502 | 8.1487 | 0 | 5 |
| 15 | Hattonville | B | 48.9924 | 5.7145 | 18 | 4 |
| 16 | Vouillon | B | 46.8257 | 1.9203 | 14 | 5 |
| 17 | Mirepoix | B | 43.158472 | 1.988042 | 3 | 0 |
| 18 | Angers | A | 47.3556 | -0.5444 | 5 | 3 |
| 18 | Angers | B | 47.3556 | -0.5444 | 0 | 1 |
| 19 | SEFRA | A | 44.8351 | 4.8911 | 16 | 0 |
| 19 | SEFRA | B | 44.8351 | 4.8911 | 4 | 0 |
| 20 | Seysses | B | 43.491 | 1.3193 | 8 | 6 |
| 21 | Moia | A | 41.8249 | 2.194 | 10 | 3 |
| 22 | San Pau | A | 42.2577 | 2.3609 | 18 | 0 |
| 23 | Col d'Ares | A | 42.3763 | 2.472 | 16 | 0 |
| 24 | Col de Jau | A | 42.6881 | 2.2509 | 1 | 0 |
| 24 | Col de Jau | B | 42.6881 | 2.2509 | 32 | 0 |
| 25 | Lusignan | B | 46.4303 | 0.1101 | 23 | 6 |
| 26 | Aigoual | B | 44.0292 | 3.4853 | 0 | 5 |
| 27 | Séranne | A | 43.7884 | 3.5433 | 12 | 0 |
| 27 | Séranne | B | 43.7884 | 3.5433 | 0 | 5 |
| 28 | Grabels | A | 43.6598 | 3.8202 | 27 | 0 |
| 28 | Grabels | B | 43.6598 | 3.8202 | 14 | 4 |
| 29 | Levernois | B | 47.0024 | 4.8472 | 5 | 5 |
| 30 | Farcheville | A | 48.4104 | 2.2889 | 11 | 4 |
| 30 | Farcheville | B | 48.4104 | 2.2889 | 8 | 3 |
| 31 | Bouleternère | A | 42.6586 | 2.5897 | 20 | 0 |
| 31 | Bouleternère | B | 42.6586 | 2.5897 | 3 | 0 |
| 32 | Versailles | A | 48.8453 | 2.0027 | 0 | 3 |

| | | | | | | |
|---|---|---|---|---|---|---|
| 32 | Versailles | B | 48.8453 | 2.0027 | 14 | 6 |
| 33 | Auxerre | B | 47.8056 | 3.6232 | 28 | 3 |
| 34 | Monterniers | A | 46.161694 | 4.623819 | 0 | 1 |
| 34 | Monterniers | B | 46.161694 | 4.623819 | 3 | 0 |
| 35 | Col de Crie | A | 46.206667 | 4.540336 | 1 | 1 |
| 35 | Col de Crie | B | 46.206667 | 4.540336 | 5 | 0 |
| 36 | Gramat | B | 44.7408 | 1.8151 | 4 | 0 |
| 37 | Sainte Baume | A | 43.3274 | 5.7176 | 6 | 4 |
| 38 | Communay | B | 45.615842 | 4.850328 | 6 | 1 |
| 39 | Maguelonne | A | 43.5098 | 3.8468 | 7 | 0 |
| 39 | Maguelonne | B | 43.5098 | 3.8468 | 13 | 1 |
| 40 | Millau | A | 44.1053 | 3.032 | 11 | 0 |
| 40 | Millau | B | 44.1053 | 3.032 | 1 | 0 |
| 41 | Montmarault | B | 46.3524 | 2.9896 | 21 | 5 |
| 42 | Lorlanges | A | 45.3361 | 3.2774 | 1 | 2 |
| 42 | Lorlanges | B | 45.3361 | 3.2774 | 10 | 0 |
| 43 | Pont du Chateaux | B | 45.803069 | 3.272247 | 8 | 0 |
| 44 | Tordera | A | 41.7725 | 2.7593 | 5 | 5 |
| 44 | Tordera | B | 41.7725 | 2.7593 | 18 | 0 |
| 45 | Firmi | B | 44.5468 | 2.3052 | 4 | 2 |
| 46 | Montagnac | A | 43.4811 | 3.4675 | 3 | 5 |
| 46 | Montagnac | B | 43.4811 | 3.4675 | 4 | 5 |
| 47 | Thézan | A | 43.1067 | 2.7447 | 3 | 6 |
| 47 | Thézan | B | 43.1067 | 2.7447 | 5 | 5 |
| 48 | Palalda | A | 42.4959 | 2.6683 | 5 | 5 |
| 49 | Boule d'Amont | A | 42.5815 | 2.6174 | 0 | 5 |
| 50 | Cuq Toulza | B | 43.5869 | 1.8487 | 9 | 5 |
| 51 | Barthe | B | 43.0786 | 0.4403 | 9 | 5 |
| 52 | Orthez | B | 43.4457 | -0.7082 | 16 | 5 |
| 53 | Saint Sever | B | 43.7809 | -0.5868 | 9 | 5 |
| 54 | Saint-Jean-Pied-de-Port | A | 43.2419 | -1.3019 | 8 | 10 |
| 54 | Saint-Jean-Pied-de-Port | B | 43.2419 | -1.3019 | 0 | 3 |
| 55 | Camargue | B | 43.0884 | 1.2881 | 8 | 5 |
| 56 | Limoux | A | 43.0559 | 2.1316 | 0 | 5 |
| 57 | Nyons | A | 44.3294 | 5.1071 | 0 | 6 |
| 58 | La Souterraine | B | 46.2279 | 1.456 | 7 | 5 |
| 59 | Ile de Ré | B | 46.187 | -1.405 | 3 | 8 |
| 60 | Moreilles | B | 46.187 | -1.405 | 1 | 1 |
| 61 | Jaulnay | A | 46.9291 | 0.4274 | 0 | 5 |
| 62 | Le Mans | A | 48.0061 | 0.3579 | 0 | 5 |
| 63 | Orleans | A | 47.8771 | 1.5647 | 3 | 9 |
| 63 | Orleans | B | 47.8771 | 1.5647 | 1 | 4 |
| 64 | Ludon | B | 44.9942 | -0.6072 | 0 | 10 |
| 65 | Velesovice | B | 49.1746 | 16.861 | 0 | 5 |
| 66 | Mrsinci | B | 43.8042 | 20.4919 | 0 | 6 |
| 67 | Villenave d'Ornon | A | 44.7764 | -0.5829 | 0 | 10 |
| 67 | Villenave d'Ornon | B | 44.7764 | -0.5829 | 0 | 1 |
| 68 | La Cavalerie | A | 44.0124 | 3.1269 | 0 | 5 |
| 69 | Guillestre | A | 44.6603 | 6.6053 | 0 | 5 |
| 70 | Magara | B | 36.7026 | 33.9083 | 0 | 25 |
| 71 | Tautavel | A | 42.7941 | 2.6985 | 0 | 5 |
| 71 | Tautavel | B | 42.7941 | 2.6985 | 0 | 12 |

| | | | | | | |
|---|---|---|---|---|---|---|
| 72 | Almese | B | 45.1167 | 7.3953 | 0 | 5 |
| 73 | Austria | B | 48.3155 | 16.3839 | 0 | 2 |
| 74 | Bosnia | B | 44.7667 | 17.1833 | 0 | 5 |
| 75 | Bursa | B | 40.1918 | 28.9854 | 0 | 10 |
| 76 | Orhaneli | B | 40.0365 | 28.8964 | 0 | 6 |
| 77 | Keles | B | 39.8957 | 29.2218 | 0 | 2 |
| 78 | Pozanti | B | 36.6273 | 33.8741 | 0 | 22 |
| 79 | Ayvali | B | 40.6206 | 42.0117 | 0 | 13 |
| 80 | Montrichard | A | 47.301442 | 1.183114 | 0 | 7 |
| 81 | Ponti | A | 44.5293 | 8.3125 | 0 | 8 |
| 82 | Loeche | A | 46.380024 | 7.628834 | 0 | 8 |
| 82 | Loeche | B | 46.380024 | 7.628834 | 0 | 8 |
| 83 | Rouen | A | 49.3045 | 1.1285 | 0 | 3 |
| 84 | Mantes la Jolie | A | 48.9861 | 1.6891 | 0 | 3 |
| 84 | Mantes la Jolie | B | 48.9861 | 1.6891 | 0 | 8 |
| 85 | Langon | A | 44.5024 | -0.2579 | 0 | 8 |
| 86 | Biarritz | A | 43.3627 | -1.5155 | 0 | 8 |
| 87 | Santander | A | 43.344 | -3.8201 | 0 | 8 |
| 88 | Quiroga | A | 42.4761 | -7.3388 | 0 | 8 |
| 89 | Venta de Banos | A | 41.9333 | -4.5016 | 0 | 8 |
| 90 | Alfaro | A | 42.1365 | -1.78 | 0 | 7 |
| 91 | Meckenheim | B | 49.4022 | 8.2186 | 0 | 8 |
| 92 | Ventiseri | B | 41.954917 | 9.382633 | 0 | 8 |
| 93 | Vescovato | B | 42.53065 | 9.4806 | 0 | 7 |
| 94 | Brest | B | 48.140198 | -4.204939 | 0 | 8 |
| 95 | Pomáz | B | 47.651035 | 19.039569 | 0 | 8 |
| 96 | Ljabojno | B | 40.89666 | 21.145213 | 0 | 8 |
| 97 | Beutenaken | B | 50.77764 | 5.856584 | 0 | 8 |

Table S2

| species (simplified name) | genre/sub-genre | species valid (after Psyl'list) | NCBI-accession | isolate | gene |
|---|---|---|---|---|---|
| Arytaina adenocarpi | Arytaina | Arytaina adenocarpi Löw, 1880 | AY100375 | Portugal | cox1-trnL-cox2 |
| Arytaina devia | Arytaina | Arytaina devia Loginova, 1976 | AY100378 | Spain | cox1-trnL-cox2 |
| Arytaina genistae | Arytaina | Arytaina genistae (Latreille, 1804) | AY100383 | United Kingdom | cox1-trnL-cox2 |
| Arytaina nubivaga | Arytaina | Arytaina nubivaga Loginova, 1976 | AY100385 | | cox1-trnL-cox2 |
| Arytaina vittata | Arytaina | Arytaina vittata Percy, 2003 | AY100386 | | cox1-trnL-cox2 |
| Arytainilla atlantica | Arytainilla | Arytainilla atlantica Percy, 2002 | AY100417 | | cox1-trnL-cox2 |
| Arytainilla cytisi | Arytainilla | Arytainilla cytisi (Puton, 1876) | AY100388 | | cox1-trnL-cox2 |
| Arytainilla delarbrei | Arytainilla | Arytainilla delarbrei (Puton, 1873) | AY100389 | | cox1-trnL-cox2 |
| Arytainilla montivaga | Arytainilla | Arytainilla montivaga Percy, 2002 | AY100419 | Spain | cox1-trnL-cox2 |
| Arytainilla serpentina | Arytainilla | Arytainilla serpentina Percy, 2003 | AY100416 | | cox1-trnL-cox2 |
| Arytainilla spartiicola | Arytainilla | Arytainilla spartiicola (Šulc, 1907) | AY100411 | | cox1-trnL-cox2 |
| Arytainilla sulci | Arytainilla | Arytainilla sulci (Vondráček, 1954) | AY100414 | | cox1-trnL-cox2 |
| Arytainilla telonicola | Arytainilla | Arytainilla telonicola Percy, 2002 | AY100418 | | cox1-trnL-cox2 |
| Arytinnis berber | Arytinnis | Arytinnis berber Percy, 2003 | AY100430 | | cox1-trnL-cox2 |
| Arytinnis canariensis | Arytinnis | Arytinnis canariensis Percy, 2003 | AY100423 | | cox1-trnL-cox2 |
| Arytinnis cognata | Arytinnis | Arytinnis cognata (Loginova, 1972) | AY100387 | | cox1-trnL-cox2 |
| Arytinnis diluta | Arytinnis | Arytinnis diluta (Loginova, 1976) | AY100391 | | cox1-trnL-cox2 |
| Arytinnis dividens | Arytinnis | Arytinnis dividens (Loginova, 1976) | AY100393 | | cox1-trnL-cox2 |
| Arytinnis equitans | Arytinnis | Arytinnis equitans (Loginova, 1976) | AY100395 | | cox1-trnL-cox2 |
| Arytinnis fortunata | Arytinnis | Arytinnis fortunata Percy, 2003 | AY100422 | | cox1-trnL-cox2 |
| Arytinnis gomerae | Arytinnis | Arytinnis gomerae Percy, 2003 | AY100428 | | cox1-trnL-cox2 |
| Arytinnis hakani | Arytinnis | Arytinnis hakani (Loginova, 1972) | AY100396 | | cox1-trnL-cox2 |
| Arytinnis hupalupa | Arytinnis | Arytinnis hupalupa Percy, 2003 | AY100429 | | cox1-trnL-cox2 |
| Arytinnis incuba | Arytinnis | Arytinnis incuba (Loginova, 1976) | AY100398 | | cox1-trnL-cox2 |
| Arytinnis menceyata | Arytinnis | Arytinnis menceyata Percy, 2003 | AY100424 | | cox1-trnL-cox2 |
| Arytinnis modica | Arytinnis | Arytinnis modica (Loginova, 1976) | AY100399 | | cox1-trnL-cox2 |
| Arytinnis nigralineata | Arytinnis | Arytinnis nigralineata (Loginova, 1976) | AY100403 | | cox1-trnL-cox2 |
| Arytinnis occidentalis | Arytinnis | Arytinnis occidentalis Percy, 2003 | AY100427 | | cox1-trnL-cox2 |
| Arytinnis ochrita | Arytinnis | Arytinnis ochrita Percy, 2003 | AY100425 | | cox1-trnL-cox2 |
| Arytinnis pileolata | Arytinnis | Arytinnis pileolata (Loginova, 1976) | AY100407 | | cox1-trnL-cox2 |
| Arytinnis proboscidea | Arytinnis | Arytinnis proboscidea (Loginova, 1976) | AY100409 | | cox1-trnL-cox2 |
| Arytinnis prognata | Arytinnis | Arytinnis prognata (Loginova, 1976) | AY100410 | | cox1-trnL-cox2 |
| Arytinnis romeria | Arytinnis | Arytinnis romeria Percy, 2003 | AY100421 | | cox1-trnL-cox2 |
| Arytinnis umbonata | Arytinnis | Arytinnis umbonata (Loginova, 1976) | AY100415 | | cox1-trnL-cox2 |
| Cacopsylla elegantula A | Hepatopsylla | Cacopsylla (Hepatopsylla) elegantula (Zetterstedt, 1840) | | NSAU0487_0102 BDV10 | cox1-trnL-cox2 |
| Cacopsylla elegantula B | Hepatopsylla | Cacopsylla (Hepatopsylla) elegantula (Zetterstedt, 1840) | | NSAU0487_0102 BDV10 | cox1-trnL-cox2 |
| Cacopsylla hippophaes A | Hepatopsylla | Cacopsylla (Hepatopsylla) hippophaes (Foerster, 1848) | | CC0542 | cox1-trnL-cox2 |
| Cacopsylla hippophaes B | Hepatopsylla | Cacopsylla (Hepatopsylla) hippophaes (Foerster, 1848) | | BDV10_NSAU0488_0101 | cox1-trnL-cox2 |
| Cacopsylla moscovita | Hepatopsylla | Cacopsylla (Hepatopsylla) moscovita (Andrianova, 1948) | | BDV10 NSAU0489_010 | cox1-trnL-cox2 |
| Cacopsylla nigrita | Hepatopsylla | Cacopsylla (Hepatopsylla) nigrita (Zetterstedt, 1828) | | BDV10_NSAU0490_0101 | cox1-trnL-cox2 |
| Cacopsylla propinqua | Hepatopsylla | Cacopsylla (Hepatopsylla) propinqua (Schaefer, 1949) | | CC0477 | cox1-trnL-cox2 |
| Cacopsylla brunneipennis | Hepatopsylla | Cacopsylla (Hepatopsylla) brunneipennis (Edwards, 1896) | | CC0478 | cox1-trnL-cox2 |
| Cacopsylla pulchra | Hepatopsylla | Cacopsylla (Hepatopsylla) pulchra (Zetterstedt, 1840) | | NSAU0494 BDV10 | cox1-trnL-cox2 |
| Cacopsylla pyri | Hepatopsylla | Cacopsylla (Hepatopsylla) pyri (Linné, 1758) | NC_038148 | | complete mitoch. genome |
| Cacopsylla pyricola | Hepatopsylla | Cacopsylla (Hepatopsylla) pyricola (Foerster, 1848) | MK039666 | | cox1-trnL-cox2 |
| Cacopsylla affinis | Thamnopsylla | Cacopsylla (Thamnopsylla) affinis (Löw, 1880) | | A109_i7 | cox1-trnL-cox2 |
| Cacopsylla alaterni A | Thamnopsylla | Cacopsylla (Thamnopsylla) alaterni (Foerster, 1848) | | gprA_A134_i5 | cox1-trnL-cox2 |
| Cacopsylla alaterni B | Thamnopsylla | Cacopsylla (Thamnopsylla) alaterni (Foerster, 1848) | | gprB_A134_i3 | cox1-trnL-cox2 |
| Cacopsylla albipes | Thamnopsylla | Cacopsylla (Thamnopsylla) albipes (Flor, 1861) | | NS0002 | cox1-trnL-cox2 |
| Cacopsylla breviantennata | Thamnopsylla | Cacopsylla (Thamnopsylla) breviantennata (Flor, 1861) | | NS0017 | cox1-trnL-cox2 |
| Cacopsylla crataegi | Thamnopsylla | Cacopsylla (Thamnopsylla) crataegi (Schrank, 1801) | | NS0005 | cox1-trnL-cox2 |
| Cacopsylla picta | Thamnopsylla | Cacopsylla (Thamnopsylla) picta (Foerster, 1848) | | A183_1i2 | cox1-trnL-cox2 |
| Cacopsylla pruni A | Thamnopsylla | Cacopsylla (Thamnopsylla) pruni (Scopoli, 1763) | MH577767 | isolate E512_13C13M | cox1-trnL-cox2 |
| Cacopsylla pruni B | Thamnopsylla | Cacopsylla (Thamnopsylla) pruni (Scopoli, 1763) | MH777774 | isolate E060_140 | cox1-trnL-cox2 |
| Cacopsylla rhamnicola | Thamnopsylla | Cacopsylla (Thamnopsylla) rhamnicola (Scott, 1876) | | CC0479 | cox1-trnL-cox2 |
| Cacopsylla albigena | Cacopsylla s.str. | Cacopsylla albigena (Miyatake, 1964) | MH769685 | | cox1-trnL-cox2 |
| Cacopsylla araliae | Cacopsylla s.str. | Cacopsylla araliae (Konovalova, 1981) | MH769688 | | cox1-trnL-cox2 |
| Cacopsylla baccatae | Cacopsylla s.str. | Cacopsylla baccatae Cho & Burckhardt, 2017 | MH769687 | isolate 45 | cox1-trnL-cox2 |
| Cacopsylla bidens | Cacopsylla s.str. | Cacopsylla bidens (Šulc, 1907) | | NS0078 | cox1-trnL-cox2 |
| Cacopsylla burckhardti | Cacopsylla s.str. | Cacopsylla burckhardti Luo, Li, Ma & Cai, 2012 | OK574466 | | complete mitoch. genome |
| Cacopsylla chinensis A | Cacopsylla s.str. | Cacopsylla chinensis (Yang & Li, 1981) | LC513974 | | cox1-trnL-cox2 |
| Cacopsylla chinensis B | Cacopsylla s.str. | Cacopsylla chinensis (Yang & Li, 1981) | AB364015 | | cox1-trnL-cox2 |
| Cacopsylla coccinea | Cacopsylla s.str. | Cacopsylla coccinea (Kuwayama, 1908) | KP245955 | | cox1-trnL-cox2 |
| Cacopsylla hakonensis | Cacopsylla s.str. | Cacopsylla hakonensis (Kuwayama, 1908) | MH769686 | | cox1-trnL-cox2 |
| Cacopsylla jukyungi | Cacopsylla s.str. | Cacopsylla jukyungi (Kwon, 1983) | LC513981 | | cox1-trnL-cox2 |
| Cacopsylla maculatili SK | Cacopsylla s.str. | Cacopsylla maculatili Li, 2011 | MK039655 | isolate 26-2 SK | cox1-trnL-cox2 |
| Cacopsylla maculatili JAP | Cacopsylla s.str. | Cacopsylla maculatili Li, 2011 | MK039657 | isolate 160-1_JAP | cox1-trnL-cox2 |
| Cacopsylla mali | Cacopsylla s.str. | Cacopsylla mali (Schmidberger, 1836) | AY100432 | | cox1-trnL-cox2 |
| Cacopsylla melanoneura | Cacopsylla s.str. | Cacopsylla melanoneura (Foerster, 1848) | | A109_6i24 | cox1-trnL-cox2 |
| Cacopsylla peregrina | Cacopsylla s.str. | Cacopsylla peregrina (Foerster, 1848) | | NSAU0344_0117m | cox1-trnL-cox2 |

| | | | | | |
|---|---|---|---|---|---|
| Cacopsylla pyrisuga | Cacopsylla s.str. | Cacopsylla pyrisuga (Foerster, 1848) | | A122_i27 | cox1-trnL-cox2 |
| Cacopsylla qianli | Cacopsylla s.str. | Cacopsylla qianli (Yang & Li, 1984) | AB364036 | | cox1-trnL-cox2 |
| Cacopsylla sandolbaea | Cacopsylla s.str. | Cacopsylla sandolbaea (Park & Lee, 1982) | MK039672 | isolate 24-2 | cox1-trnL-cox2 |
| Cyamophila floribundae | Cyamophila | Cyamophila floribundae Cho & Burckhardt, 2017 | MH769668 | isolate 128 | cox1-trnL-cox2 |
| Cyamophila prohaskai | Cyamophila | Cyamophila prohaskai (Priesner, 1927) | AY100433 | | cox1-trnL-cox2 |
| Cyamophila willieti | Cyamophila | Cyamophila willieti (Wu, 1932) | MN364946 | strain HMS1-1 | cox1-trnL-cox2 |
| Livilla adusta | Livilla | Livilla adusta (Löw, 1881) | AY100434 | | cox1-trnL-cox2 |
| Livilla baetica | Livilla | Livilla baetica Percy, 2002 | AY100443 | | cox1-trnL-cox2 |
| Livilla blandula | Livilla | Livilla blandula (Horváth, 1905) | AY100435 | | cox1-trnL-cox2 |
| Livilla caprifuga | Livilla | Livilla caprifuga Percy, 2002 | AY100442 | | cox1-trnL-cox2 |
| Livilla complexa | Livilla | Livilla complexa Percy, 2002 | AY100444 | | cox1-trnL-cox2 |
| Livilla ima | Livilla | Livilla ima (Loginova, 1972) | AY100397 | | cox1-trnL-cox2 |
| Livilla maura | Livilla | Livilla maura (Vondráček, 1951) | AY100436 | | cox1-trnL-cox2 |
| Livilla monospermae | Livilla | Livilla monospermae Hodkinson, 1990 | AY100437 | from. Spain | cox1-trnL-cox2 |
| Livilla retamae | Livilla | Livilla retamae (Puton, 1878) | AY100440 | | cox1-trnL-cox2 |
| Livilla variegata | Livilla | Livilla variegata (Löw, 1881) | AY100441 | | cox1-trnL-cox2 |
| Psylla alni | Psylla | Psylla alni (Linné, 1758) | NC038139 | | complete mitoch. genome |
| Psylla alniformosanaesuga | Psylla | Psylla alniformosanaesuga Lauterer, Yang & Fang, 1988 | MH769683 | isolate 17 | cox1-trnL-cox2 |
| Psylla foersteri | Psylla | Psylla foersteri Flor, 1861 | | CC0113 | cox1-trnL-cox2 |
| Psylla lanceolata | Psylla | Psylla lanceolata Yang, 1984 | AB364016 | | cox1-trnL-cox2 |
| Spanioneura buxi | Spanioneura | Spanioneura buxi (Linnaeus, 1758) | | NS0099 | cox1-trnL-cox2 |
| Spanioneura fonscolombii A | Spanioneura | Spanioneura fonscolombii Foerster, 1848 | | CC0041 | cox1-trnL-cox2 |
| Spanioneura fonscolombii B | Spanioneura | Spanioneura fonscolombii Foerster, 1848 | | CC0564 | cox1-trnL-cox2 |
| Spanioneura longicauda | Spanioneura | Spanioneura longicauda (Konovalova, 1986) | MH769689 | isolate 50 | cox1-trnL-cox2 |

Table S3

| Model | #Param | BIC | AICc | lnL | Invariant | Gamma | R | Freq A | Freq T |
|---|---|---|---|---|---|---|---|---|---|
| GTR+G+I | 195 | 30082,65485 | 28343,41297 | -13976,01687 | 0,390037392 | 0,69116283 | 3,91515025 | 0,348966013 | 0,399856141 |
| TN93+G+ | 192 | 30105,93978 | 28393,4346 | -14004,04872 | 0,38608061 | 0,683605561 | 3,78389238 | 0,348966013 | 0,399856141 |
| HKY+G+I | 191 | 30117,05343 | 28413,46064 | -14015,06868 | 0,400957043 | 0,759551127 | 3,36088887 | 0,348966013 | 0,399856141 |
| GTR+G | 194 | 30194,7463 | 28464,41658 | -14037,52572 | n/a | 0,281855833 | 4,34493776 | 0,348966013 | 0,399856141 |
| TN93+G | 191 | 30252,29706 | 28548,70426 | -14082,69049 | n/a | 0,315387381 | 3,27774765 | 0,348966013 | 0,399856141 |
| HKY+G | 190 | 30287,74905 | 28593,06871 | -14105,87962 | n/a | 0,321578279 | 3,13849883 | 0,348966013 | 0,399856141 |
| T92+G+I | 189 | 30373,93981 | 28688,17199 | -14154,43813 | 0,400271132 | 0,776057926 | 3,17923058 | 0,374411077 | 0,374411077 |
| T92+G | 188 | 30553,04287 | 28876,18765 | -14249,45279 | n/a | 0,323210252 | 2,8601653 | 0,374411077 | 0,374411077 |
| K2+G+I | 188 | 31346,93373 | 29670,07851 | -14646,39822 | 0,381812473 | 0,712781876 | 4,22573268 | 0,25 | 0,25 |
| K2+G | 187 | 31424,92009 | 29756,97754 | -14690,85453 | n/a | 0,340294593 | 3,8016412 | 0,25 | 0,25 |
| TN93+I | 191 | 31963,04785 | 30259,45505 | -14938,06589 | 0,435355196 | n/a | 2,54307262 | 0,348966013 | 0,399856141 |
| GTR+I | 194 | 31966,52179 | 30236,19207 | -14923,41346 | 0,438661255 | n/a | 2,74545112 | 0,348966013 | 0,399856141 |
| HKY+I | 190 | 32006,97082 | 30312,29048 | -14965,49051 | 0,438202976 | n/a | 2,6062798 | 0,348966013 | 0,399856141 |
| T92+I | 188 | 32312,31809 | 30635,46288 | -15129,0904 | 0,437470851 | n/a | 2,45948714 | 0,374411077 | 0,374411077 |
| JC+G+I | 187 | 33145,32165 | 31477,37911 | -15551,05531 | 0,391396099 | 0,86186369 | 0,5 | 0,25 | 0,25 |
| K2+I | 187 | 33149,58732 | 31481,64478 | -15553,18815 | 0,437053982 | n/a | 2,4572046 | 0,25 | 0,25 |
| JC+G | 186 | 33270,26049 | 31611,23069 | -15618,98786 | n/a | 0,362748264 | 0,5 | 0,25 | 0,25 |
| JC+I | 186 | 34926,61903 | 33267,58923 | -16447,16713 | 0,437808736 | n/a | 0,5 | 0,25 | 0,25 |
| TN93 | 190 | 35774,50144 | 34079,8211 | -16849,25582 | n/a | n/a | 2,47213934 | 0,348966013 | 0,399856141 |
| GTR | 193 | 35796,06641 | 34074,64892 | -16843,64891 | n/a | n/a | 2,21846142 | 0,348966013 | 0,399856141 |
| HKY | 189 | 35877,08719 | 34191,31938 | -16906,01182 | n/a | n/a | 2,45353156 | 0,348966013 | 0,399856141 |
| T92 | 187 | 36156,73626 | 34488,79372 | -17056,76262 | n/a | n/a | 2,45940214 | 0,374411077 | 0,374411077 |
| K2 | 186 | 36520,12152 | 34861,09172 | -17243,91838 | n/a | n/a | 2,96432193 | 0,25 | 0,25 |
| JC | 185 | 38202,89681 | 36552,77983 | -18090,76915 | n/a | n/a | 0,5 | 0,25 | 0,25 |

| Freq C | Freq G | A=>T | A=>C | A=>G | T=>A | T=>C | T=>G | C=>A |
|---|---|---|---|---|---|---|---|---|
| 0,161859378 | 0,089318468 | 0,02 | 0,02 | 0,06 | 0,02 | 0,15 | 0,01 | 0,04 |
| 0,161859378 | 0,089318468 | 0,04 | 0,01 | 0,05 | 0,03 | 0,17 | 0,01 | 0,03 |
| 0,161859378 | 0,089318468 | 0,04 | 0,02 | 0,07 | 0,03 | 0,13 | 0,01 | 0,03 |
| 0,161859378 | 0,089318468 | 0,02 | 0,02 | 0,05 | 0,02 | 0,16 | 0,01 | 0,04 |
| 0,161859378 | 0,089318468 | 0,04 | 0,02 | 0,06 | 0,03 | 0,14 | 0,01 | 0,03 |
| 0,161859378 | 0,089318468 | 0,04 | 0,02 | 0,07 | 0,03 | 0,13 | 0,01 | 0,03 |
| 0,125588923 | 0,125588923 | 0,04 | 0,01 | 0,1 | 0,04 | 0,1 | 0,01 | 0,04 |
| 0,125588923 | 0,125588923 | 0,04 | 0,01 | 0,1 | 0,04 | 0,1 | 0,01 | 0,04 |
| 0,25 | 0,25 | 0,02 | 0,02 | 0,2 | 0,02 | 0,2 | 0,02 | 0,02 |
| 0,25 | 0,25 | 0,03 | 0,03 | 0,2 | 0,03 | 0,2 | 0,03 | 0,03 |
| 0,161859378 | 0,089318468 | 0,05 | 0,02 | 0,06 | 0,04 | 0,14 | 0,01 | 0,04 |
| 0,161859378 | 0,089318468 | 0,04 | 0,02 | 0,05 | 0,03 | 0,14 | 0,01 | 0,05 |
| 0,161859378 | 0,089318468 | 0,05 | 0,02 | 0,07 | 0,04 | 0,12 | 0,01 | 0,04 |
| 0,125588923 | 0,125588923 | 0,04 | 0,01 | 0,1 | 0,04 | 0,1 | 0,01 | 0,04 |
| 0,25 | 0,25 | 0,08 | 0,08 | 0,08 | 0,08 | 0,08 | 0,08 | 0,08 |
| 0,25 | 0,25 | 0,04 | 0,04 | 0,18 | 0,04 | 0,18 | 0,04 | 0,04 |
| 0,25 | 0,25 | 0,08 | 0,08 | 0,08 | 0,08 | 0,08 | 0,08 | 0,08 |
| 0,25 | 0,25 | 0,08 | 0,08 | 0,08 | 0,08 | 0,08 | 0,08 | 0,08 |
| 0,161859378 | 0,089318468 | 0,05 | 0,02 | 0,05 | 0,04 | 0,15 | 0,01 | 0,04 |
| 0,161859378 | 0,089318468 | 0,05 | 0,03 | 0,05 | 0,04 | 0,14 | 0,01 | 0,06 |
| 0,161859378 | 0,089318468 | 0,05 | 0,02 | 0,07 | 0,04 | 0,12 | 0,01 | 0,04 |
| 0,125588923 | 0,125588923 | 0,04 | 0,01 | 0,1 | 0,04 | 0,1 | 0,01 | 0,04 |
| 0,25 | 0,25 | 0,03 | 0,03 | 0,19 | 0,03 | 0,19 | 0,03 | 0,03 |
| 0,25 | 0,25 | 0,08 | 0,08 | 0,08 | 0,08 | 0,08 | 0,08 | 0,08 |

| C=>T | C=>G | G=>A | G=>T | G=>C |
|---|---|---|---|---|
| 0,37 | 0,01 | 0,23 | 0,04 | 0,03 |
| 0,42 | 0,01 | 0,18 | 0,04 | 0,01 |
| 0,32 | 0,01 | 0,28 | 0,04 | 0,02 |
| 0,39 | 0,01 | 0,21 | 0,04 | 0,02 |
| 0,36 | 0,01 | 0,24 | 0,04 | 0,02 |
| 0,32 | 0,01 | 0,28 | 0,04 | 0,02 |
| 0,3 | 0,01 | 0,3 | 0,04 | 0,01 |
| 0,3 | 0,01 | 0,3 | 0,04 | 0,01 |
| 0,2 | 0,02 | 0,2 | 0,02 | 0,02 |
| 0,2 | 0,03 | 0,2 | 0,03 | 0,03 |
| 0,35 | 0,01 | 0,22 | 0,05 | 0,02 |
| 0,35 | 0,01 | 0,21 | 0,05 | 0,02 |
| 0,31 | 0,01 | 0,27 | 0,05 | 0,02 |
| 0,29 | 0,01 | 0,29 | 0,04 | 0,01 |
| 0,08 | 0,08 | 0,08 | 0,08 | 0,08 |
| 0,18 | 0,04 | 0,18 | 0,04 | 0,04 |
| 0,08 | 0,08 | 0,08 | 0,08 | 0,08 |
| 0,08 | 0,08 | 0,08 | 0,08 | 0,08 |
| 0,36 | 0,01 | 0,19 | 0,05 | 0,02 |
| 0,35 | 0,01 | 0,19 | 0,05 | 0,02 |
| 0,3 | 0,01 | 0,26 | 0,05 | 0,02 |
| 0,29 | 0,01 | 0,29 | 0,04 | 0,01 |
| 0,19 | 0,03 | 0,19 | 0,03 | 0,03 |
| 0,08 | 0,08 | 0,08 | 0,08 | 0,08 |

Table S4

| | |
|---|---|
| **1 sample summary statistics** | Number of haplotypes<br>Number of segregating sites<br>Mean of pairwise difference<br>Tajima's D<br>Private segregating sites<br>Mean of numbers of the rarest nucleotide at segregating sites<br>Variance of numbers of the rarest nucleotide at segregating sites |
| **2 sample summary statistics** | Number of haplotypes<br>Number of segregating sites<br>Mean of pairwise differences (W)<br>Mean of pairwise differences (B)<br>Fst (Hudson et al., 1992) |
